# How ancient forest fragmentation and riparian connectivity generate high levels of genetic diversity in a microendemic Malagasy tree

**DOI:** 10.1101/2020.11.25.394544

**Authors:** Jordi Salmona, Axel Dresen, Anicet E. Ranaivoson, Sophie Manzi, Barbara Le Pors, Cynthia Hong-Wa, Jacqueline Razanatsoa, Nicole V. Andriaholinirina, Solofonirina Rasoloharijaona, Marie-Elodie Vavitsara, Guillaume Besnard

## Abstract

Understanding landscape changes is central to predicting evolutionary trajectories and defining conservation practices. While human-driven deforestation is intense throughout Madagascar, exceptions in areas like the Loky-Manambato region (North) raise questions. Such regions also harbor a rich and endemic flora, whose evolutionary origin remains poorly understood. We assessed the genetic diversity of an endangered microendemic Malagasy olive species (Noronhia spinifolia Hong-Wa) to better understand the vegetation dynamic in the Loky-Manambato region and its influence on past evolutionary processes. We characterized 72 individuals sampled across eight forests through nuclear and mitochondrial restriction associated sequencing data (RADseq) and chloroplast microsatellites (cpSSR). Combined population and landscape genetics analyses indicate that N. spinifolia diversity is largely explained by the current forest cover, highlighting a long-standing habitat mosaic in the region. This sustains a major and long-term role of riparian corridors in maintaining connectivity across those antique mosaic-habitats, calling for the study of organismal interactions that promote gene flow.

## Introduction

One way to offset rapid anthropogenic habitat destruction and fragmentation, the primary causes of declines in global biodiversity (Fahrig, 2003; Goudie, 2018; Lindenmayer & Fischer, 2013), is to preserve habitat connectivity (Haddad et al., 2015). Such an approach is, however, insufficient by itself and requires better knowledge of species dispersal strategies (Gardner et al., 2018; Lebuhn et al., 2015; Sutherland et al., 2004), which remain poorly understood, particularly in tropical biodiversity hotspots. This typically requires understanding species diversity, their dynamic, behavior and interactions across rapidly changing landscapes (Pressey et al., 2007). Since the genetic diversity of a species is the result of both biological processes (e.g. mutation, drift, gene-flow), and effects of the environment (e.g. landscape heterogeneity), it retains the signatures of habitat and landscape changes (Balkenhol et al., 2016; Salmona, Heller, Lascoux, et al., 2017). The latter can be efficiently inferred from genetic data although assessing the relative and confounding effects of complex landscape dynamics (habitat loss, fragmentation, barriers emergence, etc.) is notoriously challenging (Beichman et al., 2018; Nater et al., 2015; Salmona, Heller, Lascoux, et al., 2017; Salmona, Heller, Quéméré, et al., 2017). Measuring the effects of landscape changes, structure, and components on genetic diversity of organisms thus remains a major issue in conservation biology.

Madagascar is considered as one of the hottest biodiversity hotspots (Myers et al., 2000). It undergoes high rates of deforestation [~40-50% area since the 1950s], resulting in habitats loss and fragmentation, and ultimately biodiversity loss (Harper et al., 2007; Vieilledent et al., 2018). Anthropogenic deforestations since human settlement on the island have long been assumed to have generated the Malagasy grasslands. However, their Miocene origin (Bond et al., 2008; Hackel et al., 2018; Salmona et al., 2020; Solofondranohatra et al., 2018; Vorontsova et al., 2016) points to the antiquity of Madagascar’s habitat mosaics, fueling a hot debate on the causes and age of its habitat fragmentation (Godfrey & Crowley, 2016; Joseph & Seymour, 2020, 2021). Meanwhile, it is a place where connectivity preservation would be a very impactful conservation approach (Morelli et al., 2020). Yet, an understanding of the nature of the fragmentation, whether natural or human-induced and/or ancient or recent, is essential to better assess how species are molded by landscape changes, and to consequently factor their survival strategies in an effective conservation plan.

The Loky-Manambato (LM) region in northern Madagascar, which features perplexingly mild deforestation (Quéméré et al., 2012; Salmona, Heller, Quéméré, et al., 2017), shows well-characterized matrix of forests and open-habitats, a diversity of putative barriers to gene flow, as well as high levels of endemism across living kingdoms (Goodman et al., 2018; Goodman & Wilmé, 2006). It rose as a small-scale model-region to investigate the effects of landscape changes, structure, and components on species’ genetic diversity. For instance, the forest-matrix was identified among the major landscape components shaping genetic diversity across all species studied in the LM region (Aleixo-Pais et al., 2019; Quéméré et al., 2010; Rakotoarisoa et al., 2013a; Sgarlata et al., 2018; Tang et al., 2020). Contrastingly the Manankolana River, showed a dominant effect on *Propithecus tattersalli* Simmons, not recovered in other species. Aside mammals, contributions on other taxa, such as plants, are crucial to draw taxonomically-broad generalities regarding the history, connectiveness, and conservation value of the current LM landscape.

By their very nature, native tree species are at the forefront of deforestation and landscape changes. Trees, therefore, represent putatively good models for landscape genetics studies in fragmented habitats, despite their long generation time. However, only a few studies have used the genetic diversity of Malagasy plant populations (Andrianoelina et al., 2009; Gardiner et al., 2017; Salmona et al., 2020) to infer landscape dynamics and inform conservation strategies. The Malagasy olives (genus *Noronhia* Stadt. ex Thouars), with a high number of taxa and a high microendemism rate, are among the major components of Madagascar forests and of the LM region in particular (Hong-Wa, 2016; Hong-Wa & Besnard, 2014). Among them, the Malagasy spiny olive (*Noronhia spinifolia* Hong-Wa) is mostly endemic to the dry to sub-humid forests of the LM region and is of high conservation concern due to its narrow range. Nonetheless, *N. spinifolia* is quasi-ubiquitous in the LM region such that its genetic makeup is expected to reflect the signals of the processes operating on its populations across time and space. Moreover, being narrowly distributed, it may hold relatively low genetic diversity (Kimura, 1983) and suffer from inbreeding depression due to recent population collapse. Although its pollen and seed dispersal have yet to be studied, *N. spinifolia’s* flower and fruit morphology suggests insect pollination and animal-mediated dispersal of fruits (see below). *Noronhia spinifolia*, therefore, represents an excellent model to better understand Malagasy olives’ ecology and offers a case study to define appropriate action for dry-forests plant conservation in northern Madagascar.

In such sexually-reproducing plants, dispersal occurs by two means: via haploid male gametes in pollen, and via diploid embryos in seeds. Without field data, population and landscape genetics offer an alternative way to estimate effective dispersal (Balkenhol et al., 2016; Holderegger et al., 2010). In particular, the combined use of complementary maternally and biparentally inherited genetic data [respectively from chloroplast or mitochondrial genomes (cpDNA or mtDNA) and the nuclear genome (nDNA)] allows disentangling, to a certain level, the relative contribution of seed and pollen dispersals in gene flow (Cruzan & Hendrickson, 2020). For instance, the congeneric *N. lowryi* Hong-Wa exhibited contrasting near-panmixia nuclear and strong chloroplast genetic structures suggesting a long and short distance dispersal of pollen and seeds, respectively (Salmona et al., 2020). While progresses in sequencing technologies facilitated the generation of such genetic data for non-model organisms (Allendorf et al., 2010), recent advances in spatially explicit analyses also unlocked our ability to estimate the effect of numerous collinear landscape features on genetic diversity (Balkenhol et al., 2016; Prunier et al., 2017). Furthermore, although the limited number of tested alternative landscape hypotheses long relied on prior knowledge or expert opinions, recent approaches iterating around a large panel of resistance values (Graves et al., 2013) or optimizing resistance surfaces (Peterman, 2018; Peterman & Pope, 2021), widened the potential to identifying relevant landscape components while optimizing their cost values from the genetic data itself.

The overall goal of our study is to assess the effects of landscape on *N. spinifolia*’s genetic diversity and population structure to inform conservation strategies. We used genomic data from recently collected specimens of *N. spinifolia* across most of its range, the LM region. We first tested whether its restricted geographic distribution resulted in a low genetic diversity, as expected under a neutral model (Kimura, 1983), or remained relatively high as for co-distributed primates *[P. tattersalli* and *Microcebus tavaratra* Rasoloarison, Goodman & Ganzhorn (Aleixo-Pais et al., 2019; Quéméré et al., 2010)]. We then measured the effect of landscape components on biparentally and maternally inherited genetic diversity, to investigate patterns of pollen and seed dispersals. We compared its dispersal patterns with those of a congeneric species from the High Plateau [*N. lowryi* (Salmona *et al*., 2020)] that exhibits near-panmixia on pollen-dispersed genes, but very short seed dispersal, and with those of co-distributed mammal taxa (abovementioned), whose dispersal is mainly limited by non-forest habitats and rivers. The little knowledge about its pollen and seed dispersal agents does not allow making strong predictions, except that dispersal will depend on the vectors and on their use of the landscape. We also examined whether the relative stability of the forest cover in the past 70 years (Quéméré et al., 2012) is reflected in *N. spinifolia* genetic makeup, comparing the effect of recent and historical forest covers on gene flow, as a proxy for the temporality of its habitat loss and fragmentation. Finally, we present the application of our work to the conservation of the LM region forest network.

## Material and methods

### Study region

The Loky-Manambato (LM) region (Daraina; Fig. 1) is a biogeographical transition zone between dry deciduous and humid forests (Goodman & Wilmé, 2006), stretched over ~2,500km^2^ between the Loky and Manambato Rivers. This region is crossed by the relatively shallow Manankolana River, bordered by riparian forests along most of its course, and by a national dirt road (Fig. 1). It consists of a dozen major forest patches, ranging from 8.3 to 62.5 km^2^ and totaling ~360 km^2^ (Goodman et al., 2018), separated by ~0.5-10 km of human-altered grasslands, dry scrub, agricultural lands and riparian corridors. Most forests are situated at low-to mid-elevations and mostly consist of dry deciduous vegetation. In contrast, some mountain forests (Binara and Antsahabe, plus Bobankora to a lower extent) are covered by a gradient of dry deciduous, transition, humid and ericoid vegetation (Gautier et al., 2006). Despite sustained grassland fires, slash-and-burn agriculture and charcoal production, as well as exploitation of wood, gold and sapphires (Fanamby, 2010; Goodman *et al*., 2018), deforestation rate in the LM region is still relatively low (Quéméré *et al*., 2012) compared with those of eastern and southwestern Madagascar (Vieilledent et al., 2018), likely stemming from its remoteness, difficult accessibility and climate. However, to mitigate the threats, the LM region progressively became managed as a protected area by the Malagasy NGO “Fanamby” since 2005 (Fanamby, 2010; Goodman *et al*., 2018).

**Figure 1:**
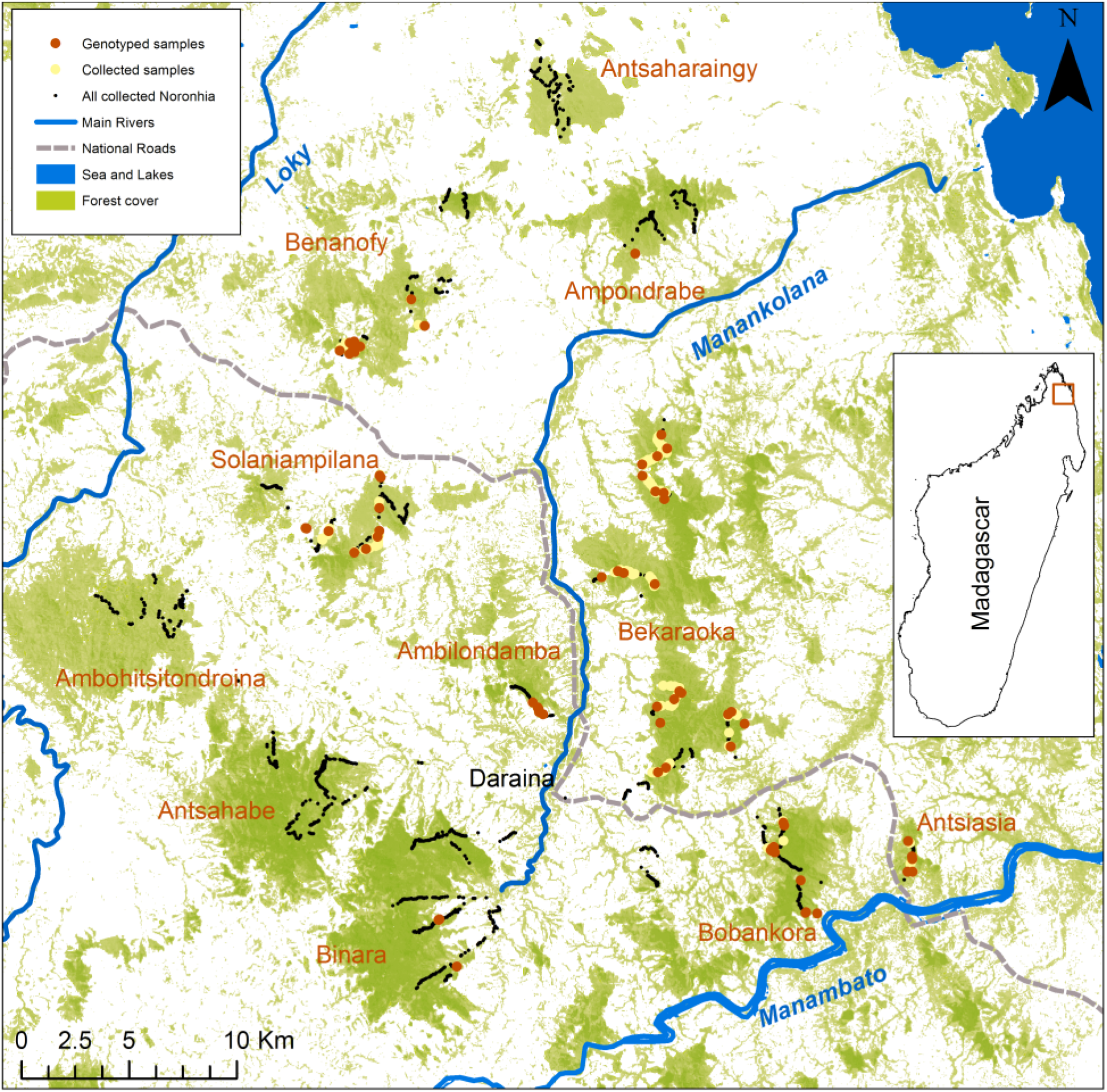
Map of *Noronhia spinifolia* sampling in the Loky-Manambato (LM) region. The small black points represent samples collected for all *Noronhia* species (ca. 30 distinct taxa) and illustrate the survey effort conducted in the region. The yellow and red dots represent *N. spinifolia* samples, with the red dots corresponding to samples included in our genomic analyses. The forest cover is adapted from Hansen *et al*. (2013). Pixels with less than 30% tree cover are represented in white to illustrate the presence of riparian forests along streams of the LM region.

### Study species

*Noronhia spinifolia* (Oleaceae) is a small-sized, understory tree that is easily distinguished from other *Noronhia* species by its narrow linear leaves with a spiny tip. The plant has cream-white, urceolate, small (< 7 mm long), and hermaphroditic flowers, as well as small (<10 mm long) and drupaceous fruits that have a thin mesocarp and a rather crustaceous endocarp. Flowering and fruiting typically occur from October to May, during the rainy season. Flower and fruit characteristics, along with observational accounts, suggest insect pollination [e.g. bees] and animal dispersal [e.g. birds, lemurs, rodents] (Hong-Wa, 2016), which may be short-distanced depending on the nature and behavior of the vectors (e.g. forest-dwellers). It is microendemic to northern Madagascar, mainly found in the LM region except for one record from further north in Montagne des Français, and is reported mainly in semi-deciduous forests of low elevation, mostly on alkaline substrate [e.g. limestone, calc-alkaline rocks]. *Noronhia spinifolia* has been assigned a preliminary conservation status of “Endangered” due to threats to its habitat (Hong-Wa, 2016).

### Plant sampling

To sample *N. spinifolia* populations, we surveyed all major forests of the LM region (Fig. 1) in 2017 and 2018, during the dry season (July-September), and used topography (altitude and shape) as a sampling guide to maximize the representation of all landscape features. Most surveys (black dots in Fig. 1) started from the forest edge at low altitude towards the forest core at higher elevation. We identified *Noronhia* species based on tree characteristics, leaf morphology and tissue structure, and collected leaf samples of 220 *N. spinifolia* trees - which tends to occur in patches - preserved in silica gel for DNA conservation. We prioritized fully-grown mature tree sampling because much of the density-dependent mortality takes place before maturity in trees, and their effective population size contributing to the genetic diversity is thus closer to the actual adult census size than to the size of the entire population including young trees and seedlings (Dodd et al., 1999; Petit & Hampe, 2006). Therefore, the regional patterns of diversity are expected to be better represented by adult samples. For each tree, we systematically recorded its height, diameter and reproductive state, as well as its geographical coordinates (GPS) and elevation. For all forests, at least one specimen voucher was prepared and deposited at the herbarium of the Parc Botanique et Zoologique de Tsimbazaza (TAN).

### Laboratory procedures

#### DNA extraction, organellar and nuclear genotyping

We extracted DNA from 137 samples of *N. spinifolia* using the Qiagen Biosprint DNA Plant kit, followed by quality control procedures ensuring high quality genomic DNA. We subsequently genotyped 72 high DNA quality samples (Fig. 1, Methods S1); a cost-effective subsampling that nonetheless maximizes geographic and altitudinal representation, tends to limit clustered sampling, and retains samples from all geographic distances, and that also prioritizes reproductively mature and fully-grown trees with a targeted sequencing depth >15×. Using a two-pronged approach, we genotyped 15 chloroplast microsatellites (cpSSR) and one mitochondrial microsatellite (mtSSR), originally developed on *Olea europaea* L. (Table S1, Methods S2, S3; Besnard *et al*., 2011), and also used restriction associated DNA sequencing (RADseq; generating data from the biparentally inherited nuclear genome and the mitogenome; Methods S4). RADseq consists in sequencing regions neighboring restriction sites, to obtain homologous sequences across individuals, spread across the genome, at a reasonable cost (Andrews et al., 2016; Baird et al., 2008).

### Data processing

#### Organellar loci, nuclear ploidy and de-novo assembly of the nuclear loci catalog

After ad-hoc demultiplexing and cleaning of reads (Methods S4), we screened the organellar genomes using bwa-mem sequence alignment (Li, 2013) to the *N. clarinerva* Hong-Wa mitogenome and *N. spinifolia* plastome (MW202230 and MT081057, respectively; Methods S5). We identified ten mitochondrial *Sbf*I RAD (mtRAD) loci *in silico*, from which haplotypes were called using ANGSD v0.92 (Korneliussen et al., 2014; Nielsen et al., 2012), based on their highest effective base depth (Wang et al., 2013). Conversely, no cpDNA RAD locus was recovered, confirming *in silico* analyses (Methods S5). Organellar genomes are supposedly inherited from the mother in the olive tribe, and consequently their polymorphism should be closely linked (Besnard et al., 2002; Van de Paer et al., 2018). As we cannot directly verify this assumption on progenies of *N. spinifolia*, we here investigated the linkage disequilibrium [LD] among organellar markers using the package *poppr* [Method S2 (Agapow & Burt, 2001; Kamvar et al., 2014, 2015)].

A catalog of nuclear tags (loci) was *de-novo* optimized (Methods S6) by iterating around the core parameters of Stacks (Rochette et al., 2019) to maximize the amount of available biological information (Paris et al., 2017). The final catalog was further cleaned (Methods S6) for exogenous contaminants using DeconSeq (Schmieder & Edwards, 2011) and endogenous orthologs using MUMmer (Kurtz et al., 2004). To enable ploidy-adapted analyses (catalog assembly and variants genotyping), ploidy was inspected using minor allele frequency plots and statistically confirmed using nQuire (Weiß et al., 2018).

#### RADseq genotyping

We used two fundamentally distinct genotyping approaches to ensure the robustness of our results: single nucleotide polymorphism (SNPs) called in Stacks, and genotype likelihoods (GLs) estimated with ANGSD (Methods S7). GLs retain information about uncertainty in base calls, which alleviates some issues associated with RADseq data such as unevenness in sequencing depth and allele drop-outs (Heller et al., 2021; Pedersen et al., 2018; Warmuth & Ellegren, 2019).

### Population and landscape genetics

We conducted complementary analyses to assess the effect of landscape components on the genetic diversity of *N. spinifolia*. We first investigated the raw patterns of genetic diversity and structure without priors to describe the major trends and build hypotheses. Then, using univariate approaches under an isolation-by-resistance model (IBR; McRae, 2006), we assessed the effect of each landscape component, iterating through their cost (resistance or conductance) and resolution (i.e. granularity or pixel size). Finally, we optimized the composite landscape resistance surface using a gradient based approach to assess the relative contribution of selected landscape components to *N. spinifolia*’ s genetic connectivity across the area.

#### Genetic diversity

To assess *N. spinifolia’*s genetic diversity, its distribution in space and among genomic compartments, we estimated several diversity metrics. We estimated both the individual and the within-forest-averaged expected heterozygosity *(H_E_)* according to Fumagalli (2013), as well as the proportion of heterozygous genotypes (*H*_O_), from nuclear genotype likelihoods (GL), based on unfolded site frequency spectra estimated in ANGSD. We further estimated organellar diversity (*h*), the probability that two haplotypes are different (Nei, 1987).

#### Population structure

To understand how *N. spinifolia*’s population is structured across the LM region, we first used forest-based metrics and naive multivariate approaches. We assessed the level of genetic differentiation among forests with Reynolds’ weighted *F_ST_* (Reynolds et al., 1983) from GL inferred in ANGSD. We explored the genetic structure of our study system through naive clustering analyses (Methods S8), based on ANGSD GLs using NgsAdmix v32 (Skotte et al., 2013) and on Stacks called genotypes using ADMIXTURE v1.3.0 (Alexander et al., 2009), and with a principal component analysis (PCA) from GLs with PCAngsd. We estimated the level of organellar genetic differentiation among forests with Nei’s weighted *F*_ST_ (Nei, 1973), Nei’s *D* (Nei, 1973), and Edwards’ chord distance [Edwards’ D (Edwards, 1971)], based on frequency of haplotypes found in each forest and appropriate for organellar material haplotypic data (Grasty et al., 2020; Kohrn et al., 2017), using the R packages *hierfstat* and *adegenet*. We also investigated the phylogenetic structure of organellar DNA data using minimum spanning networks of genetic distances (see below) constructed with the R package *poppr* (Kamvar et al., 2015).

#### Isolation by distance

To assess how the geographic distance alone explains the above-mentioned among-individual genetic relationships [distance or relatedness], we investigated patterns of isolation by distance [IBD (Slatkin, 1993; Wright, 1943)]. We used Mantel tests (Mantel, 1967) between individual geographic and genetic distances (Methods S9). IBD may be limited to a certain scale and disrupted at larger scales (e.g. Keller & Holderegger, 2013; Van Strien *et al*., 2015; Cayuela *et al*., 2019), due to effective dispersal over recent or longer evolutionary time frames (Cruzan & Hendrickson, 2020). To assess the scale at which IBD is maximized, we compared data subsets upper-bonded by their maximum geographic distance (S) (Methods S9), and which represent the cumulative effect of all data included below S. In addition, to evaluate how effective dispersal is distributed geographically, we build Mantel correlograms (Oden & Sokal, 1986; Sokal, 1986), using the *ecodist* R package (Goslee & Urban, 2007).

We used the Bruvo’s genetic distances (Bruvo et al., 2004) to consider the specific mutation model of the cpSSR data. Additionally, we used Nei’s genetic distance [Nei’s *D* (Nei, 1973)], appropriate for small scale study of organellar material haplotypic data [(Kohrn et al., 2017) Methods S3]. Finally, we estimated the covariance of nuclear RADseq GLs (Meisner & Albrechtsen, 2018), in PCAnsgd, as such metrics based on principal components analysis have been shown to be particularly fitted for landscape genetics (Shirk et al., 2017).

#### Isolation by resistance

Landscapes are rarely homogeneous, and gene flow may be limited or facilitated by its components. To assess the cost (i.e. resistance/conductance) associated with effective dispersal through each landscape feature, we used an IBR approach (McRae, 2006). As detailed below, we assessed the unique and common contributions of each landscape variable, at varying cost and resolution (i.e. granularity or pixel resolution), for two movement models [Least Cost Path (Adriaensen et al., 2003) and Circuit Theory (McRae & Beier, 2007)], to the individual genetic distances of *N. spinifolia*, for each of the three genetic compartments (cpDNA, mtDNA, nDNA).

#### Landscape variables, cost and resolution

As *N. spinifolia* was recently described and occurs in a remote area (Hong-Wa, 2016), we had little prior knowledge on the landscape variables that may affect pollen and seed dispersal. We therefore assessed the effect of most available landscape variables (Table 1; Methods S10). To test if the genetic diversity of old trees may be better explained by past forest cover, we used forest cover data from 1953, 1973, and 2000s (Hansen et al., 2013; Vieilledent et al., 2018). Although strong priors associating a landscape component to a particular cost (resistance or conductance) may be available for well-studied species (e.g. Dellicour *et al*., 2019; Quéméré *et al*., 2010), landscape variables and their associated cost are often chosen almost arbitrarily when little or no data are available (Beier et al., 2008, 2011). To identify the variable-cost associations that matter for our study system, we iteratively tested 14 conductance-resistance values (Methods S10). Similarly, organisms do not necessarily perceive each environmental component at the same resolution [or granularity, expressed as pixel size (Chubaty et al., 2020; Galpern et al., 2012; Galpern & Manseau, 2013)]. To identify the variable-cost-granularity relevant for *N. spinifolia*, we tested four pixel resolutions (Methods S10).

**Table 1:**
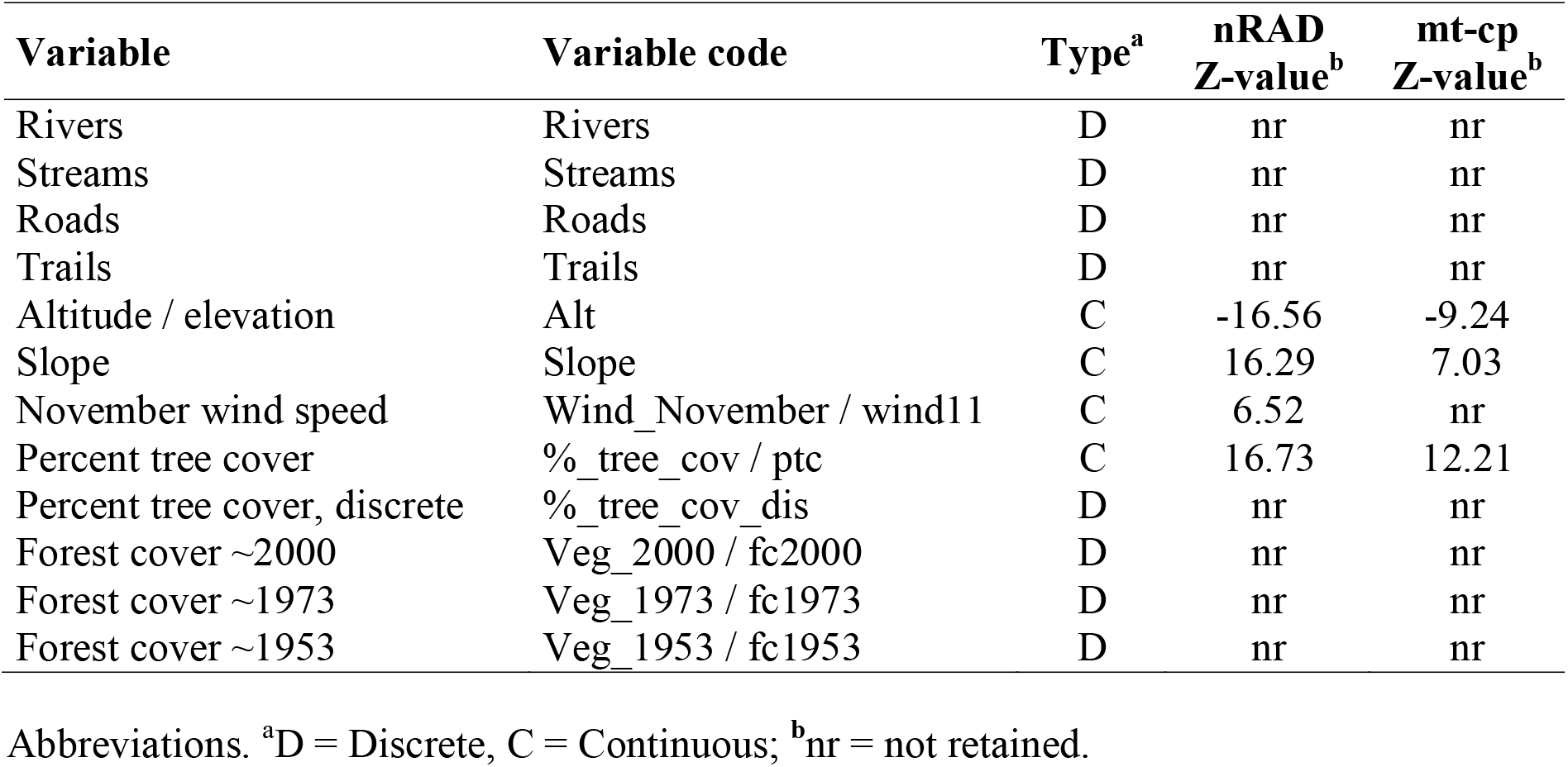
Landscape variables.

#### Movement model and analytical flow

To determine which dispersal model best applies to *N. spinifolia*, we used both the Least Cost Path (LCP) and the Circuit Theory (CT). These two approaches, respectively, consider the least cost trajectory and the cost of all possible trajectories (McRae & Beier, 2007). We computed landscape distances using the R package *gdistance* (van Etten, 2012).

We used a two-step procedure to assess the landscape components’ relative contributions to the spatial structure of *N. spinifolia’* s genetic diversity. First we selected landscape components, as well as their best fitting cost, resolution, and movement model. We estimated the correlation between geographic or landscape distance and genetic matrices (i.e. Landscape variables and Genetic distances as described above) using Mantel tests (Mantel, 1967) in the R Package *vegan* (Dixon, 2003). We retained variables showing a better fit *(R^2^)* than IBD, exhibiting sensitivity to cost values (i.e. variables with a fixed fit across all cost values were discarded), and investigated their sensitivity to cost, movement model, and resolution. Second, we optimized the composite landscape conductance surface using the gradient based approach implemented in *radish* (Peterman & Pope, 2021) to assess the contribution of landscape variables, using a log linear conductance and a maximum likelihood population-effect parameterization of a mixed effects model [MLPE (Clarke et al., 2002; van Strien et al., 2012)]. MLPE accommodates the nonindependence of pairwise genetic and environmental distance and was assessed as the most accurate regression-based approach when conducting landscape genetics model selection in individual-based analyses (Shirk et al., 2018). Because the first and second step models are relatively distinct, we assessed the contribution of the variables retained from the first procedure step, as well as variables required to test specific dispersion hypotheses (wind, slope, streams network, etc.). To limit the analytical computation load, we hierarchically build models of increasing complexity (with one to four variables) combining the variables showing the best fit (AIC), and a significant effect at the previous hierarchical level. Altogether, we compared 32 organellar and 59 nuclear optimized composite landscape conductance models (Tables S2 and S3).

## Results

### Species occurrence

We sampled *N. spinifolia* in eight of the 11 surveyed major forests of the LM region (Fig. 1). The species occurs from low to medium elevation, between 87 and 505 m, but with strong discrepancies among forests (Fig. S1). While it was mainly recorded in dry forests, it was surprisingly found in dry to wet transition forests at medium elevation (451-505 m) in Binara. Furthermore, the species was not found in three major forest patches of the LM region - namely Antsahabe, Ambohitsitondroina and Antsaharaingy - despite (*i*) large prospection efforts in these forests, and (*ii*) apparently similar habitat as the neighboring forests harboring the species (Fig. 1).

### Organellar genotyping, nuclear ploidy and catalog construction

Of the 15 chloroplast microsatellites, 14 showed polymorphism (Table S4), and allowed distinguishing 55 chlorotype profiles among 72 trees (Results S1). The ten mitochondrial RAD loci (mtRAD) allowed identifying 11 SNPs (Results S1; Table S5). The combination of mtRADs and the mtSSR locus permits the identification of 15 mitotypes among 72 trees (Table 2). Contrasting LD patterns were found within and among organellar markers (Fig. S2; Results S1), suggesting the occurrence of recombination in the mitogenome possibly due to occasional paternal leaks of mitochondria.

**Table 2:** Chloroplast, mitochondrial and nuclear summary statistics.

| Forests | cpSSR |  |  |  | mtRAD |  |  |  | nRAD |  |
| --- | --- | --- | --- | --- | --- | --- | --- | --- | --- | --- |
| | N | $n_h$ | $H$ | $A_r$ | N | $n_h$ | $H$ | $A_r$ | N | $H_E$ |
| Ambilondamba | 6 | 5 | 0.98 | 2.22 | 5 | 4 | 0.97 | 2.16 | 5 | 0.00510 |
| Ampondrabe | 1 | 1 | - | - | 1 | 1 | - | - | 1 | - |
| Antsiasia | 6 | 4 | 0.92 | 2.67 | 6 | 3 | 0.81 | 2.14 | 6 | 0.00563 |
| Bekaraoka | 25 | 19 | 0.99 | 2.38 | 23 | 5 | 0.66 | 2.04 | 23 | 0.00652 |
| Benanofy | 11 | 8 | 0.94 | 2.39 | 11 | 4 | 0.78 | 2.26 | 11 | 0.00559 |
| Binara | 5 | 2 | 0.73 | 2.36 | 5 | 2 | 0.73 | 2.05 | 5 | 0.00453 |
| Bobankora | 11 | 10 | 0.99 | 2.45 | 11 | 3 | 0.73 | 2.04 | 11 | 0.00624 |
| Solaniampilana | 10 | 8 | 0.97 | 2.37 | 10 | 5 | 0.87 | 2.17 | 10 | 0.00624 |
| Total / Mean | 75 | 55 | 0.99 | 2.41 | 72 | 15 | 0.85 | 2.12 | 72 | 0.00676 |

N = number of analyzed individuals; $n_h$ = number of haplotypes; $h$ = haplotype diversity; $A_r$ : allelic richness (estimated for five individuals), $H_E$ = expected heterozygosity.

Individual-based minor allele frequency profiles displayed unimodal diploid patterns, confirmed by nQuire analyses, and echoing the low frequency of polyploid in the genus *Noronhia*(Gorrilliot et al., 2021). The nuclear catalog parameter space exploration iterating around the core parameters for Stacks [i.e. *m* – the minimum number of reads required to build a stack, *M* – the maximum number of differences between stacks of an individual allowed when building a locus; and *N* – the maximum number of differences between loci of multiple individuals allowed when building a loci] allowed selecting values (*m* = 4, *M* = 5, *N* = 8) that offer a trade-off between the coverage, loci number, and SNP number, while limiting the number of paralogs and the presence of contaminants (Results S2). The SNP-calling procedure showing low ability to recover the genetic makeup of *N. spinifolia* (when compared to the GL-based procedure; Figs S3-S8), we therefore limited its use to preliminary analyses (ADMIXTURE & genetic distances) and proceeded with the GL-based procedure for downstream analyses.

### Genetic diversity

Chloroplast microsatellites revealed a relatively high genetic diversity with only two chlorotypes shared by individuals from more than one forest, resulting in a high probability that two randomly sampled haplotypes are different (*h* = 0.99) and a mean allelic richness (*A_r_*; estimated for five individuals) of 2.41 (Table 2). Consequently, most forests showed an extremely high cpSSR genetic diversity (*h* > 0.92) with the exception of Binara that appeared slightly less diverse (*h* = 0.73; Table 2). A relatively high mitotype diversity was also revealed [*h* = 0.85 (ranging from 0.66 to 0.97 per forest), *A_r_* = 2.12]. Contrastingly, most forest patches exhibit moderately high levels of nuclear diversity with *H_E_* values ranging from 4.53 to 6.52 × 10^-3^, with discrepancies within and among forests (Tables 2 and S1; Fig. S9). This diversity is not homogeneously distributed in space, and higher levels of genetic diversity seemingly occur in certain areas such as Solaniampilana (Fig. S10). Furthermore, genetic diversity does not seem influenced by altitude (Fig. S11).

### Population structure

The chloroplast and mitochondrial data both revealed substantial differentiation among forests (*F*_ST_ estimates ranging from 0.040 to 0.393 for cpSSRs; and 0.005 to 0.661 for mtRADs). As expected, a strong differentiation was also observed when combining cpDNA and mtDNA data (*F*_ST_ estimates ranging from 0.101 to 0.401; Table S6). The Solaniampilana and Benanofy forests, the farthest apart from other forest patches, formed an easily distinguishable haplotype group for both mtDNA and cpDNA (Figs S12-S13), while other more adjacent forests showed more limited divergence among them. Haplotype networks based on cpSSR and/or mtRAD data also revealed that one maternal lineage is unique to Solaniampilana and Benanofy (Fig. 2). Furthermore, the geographic Euclidean distances showed low, but highly significant, power at explaining genetic distances among individuals (*R*^2^ [cpSSR]: 11.7%; *R*^2^ [mtRAD]: 20.7%; and *R*^2^ [cpSSR + mtRAD]: 21.3%; Figs S8, S14, S15; Results S3) and supported by the Mantel correlogram positive and significant values within 10-12 km classes (Fig. S16).

**Figure 2:**
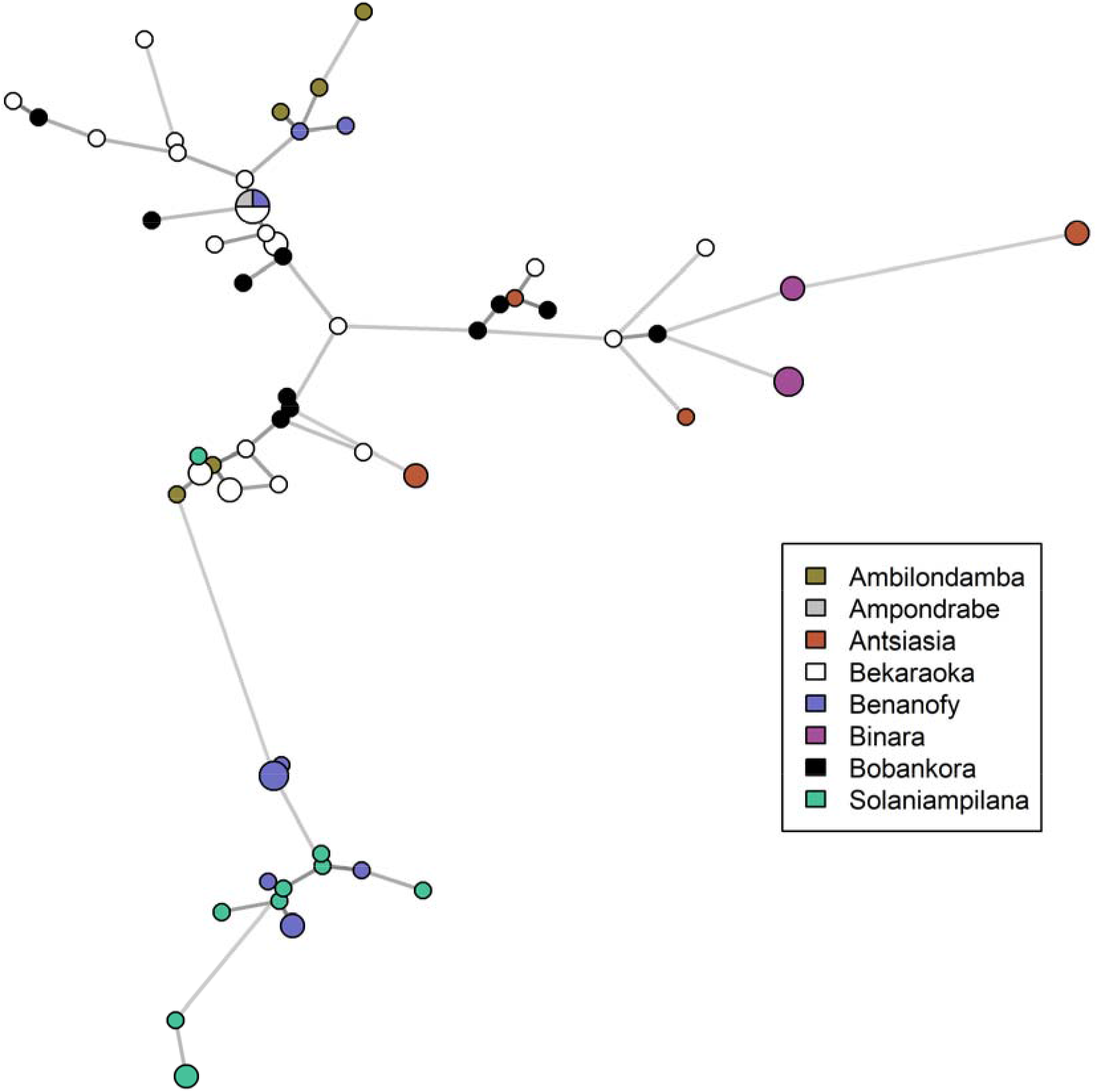
Organellar DNA haplotype network of *Noronhia spinifolia*. Line length and gray scale are proportional to the Bruvo’s cpDNA + Manhattan mtDNA combined genetic distances between distinct organellar haplotypes. Pie chart size is proportional to the occurrence number of a given haplotype. All edges of equal weight are represented. Distances among haplotypes are represented both through longer edges and the gray scale. The network highlights the huge organellar DNA diversity in *N. spinifolia*, with only one haplotype shared by individuals from at least two forests. It further shows a limited spatial structure, with, for instance, haplotypes from Solaniampilana and Benanofy grouping together at the bottom of the network.

*F_ST_* estimates based on nuclear markers (Table S7) ranged from 0.089 to 0.210, indicating that most forests are differentiated from each other. However, we found no strong structure in subpopulations, with no particular support for number of clusters >1, both for GL- and SNP-based analyses (Figs S3, S4). Instead, we found a clear northwest-southeast signal of continuous genetic differentiation across space, through GL-based PCA (First axis, ~15% of the variance explained; Fig. S17), clustering (Figs 3, S5, S6), and IBD analyses (Figs S8, S15). The observed continuous structure is well illustrated by the clustering structure for *K* = 3 that shows admixed patterns at all sampling sites (Fig. 3). We found a clear IBD signal explaining up to 56.6% of the among-individuals nuclear GL covariance (Fig. S15), further supported by the Mantel correlogram positive and significant values within 14 km classes (Fig. S16).

**Figure 3:**
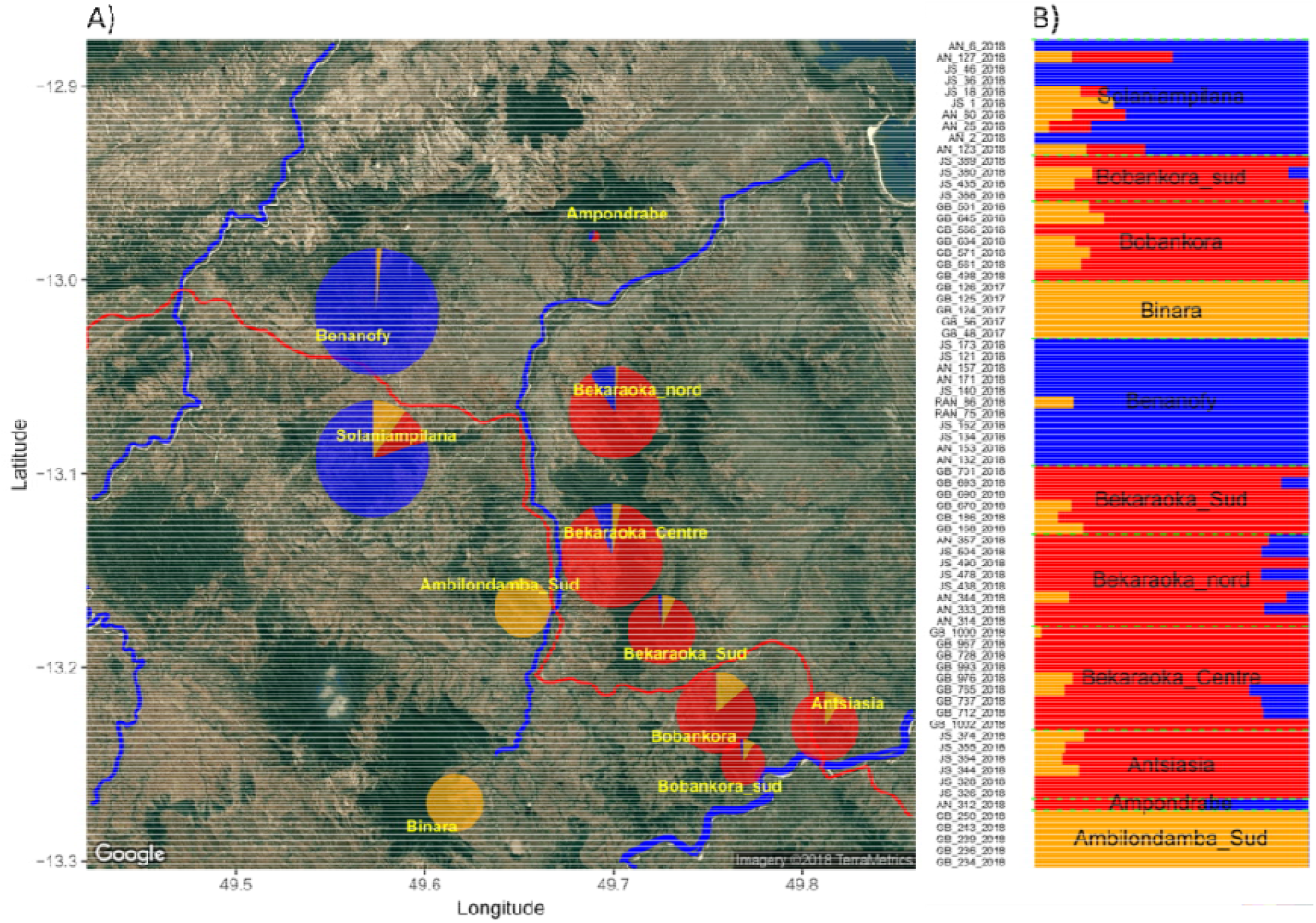
Spatial genetic structure of *Noronhia spinifolia* in the Loky-Manambato region. NgsAdmix ancestry proportions (for *K* = 3 genetic clusters) represented either **(A)** spatially by sampling site, or **(B)** per individual. Size of pie charts (in A) is proportional to the number of samples per site. Pie shares represent the sums of individual ancestry proportions that are shown in B. Results are arbitrarily represented for *K* = 3, according to the likelihood and delta*K* results in Fig. S8, because this *K* value best illustrates the continuous pattern of structure inferred using ngsAdmix and other approaches.

### Landscape genetics

The univariate Mantel tests of genetic and landscape distances (IBR) showed lower linear fit for cpDNA *(R^2^* max ~0.14) than for mtDNA (*R*^2^ max ~0.38) and nDNA (*R*^2^ max ~0.90). Among the four vegetation layers, the continuous and discrete percent tree cover layer always exhibited the highest fit for conductance values at high resolution with cpDNA, mtDNA and nDNA individual-based genetic distances *(R^2^* = 0.14; 0.38 and 0.90, respectively; Figs S18-S21). Altogether, the parameter space exploration reveals a strong effect of all forest cover layers, whereas some other variables (i.e., rivers, roads and slope) may have subtle lower effects too. To build multivariate models, we retained in priority landscape variables showing a better fit (*R*^2^) than the null model (IBD), and exhibiting sensitivity to cost values (e.g. % forest cover).

The hierarchically-built composite landscape conductance surface of increasing complexity confirmed that recent forest cover layers – the percent of tree cover, and the forest cover in 1973 to a lower extant – showed high univariate fit and log-collinearity with nuclear and organellar genetic data (Fig. S22). In addition, among monthly wind speed, that of November showed the best fit and highest collinearity with nuclear data (Fig. S23). The best fitting composite landscape conductance surface optimization for nuclear data (Table S3) retained the percent of tree cover (Hansen et al., 2013), the elevation (altitude), the wind speed in November and the slope (Table 1, Fig. 4). Similarly, the model best fitting the organellar data (Table S2) retained the percent of tree cover, the elevation and the slope (Fig. S24). For both nuclear and organellar data, the percent of tree cover always displayed the highest positive z-value for all covariates (Table 1), whereas the elevation was the only variable exhibiting negative relationship with genetic distances (Table 1, Figs 4 and S24).

**Figure 4:**
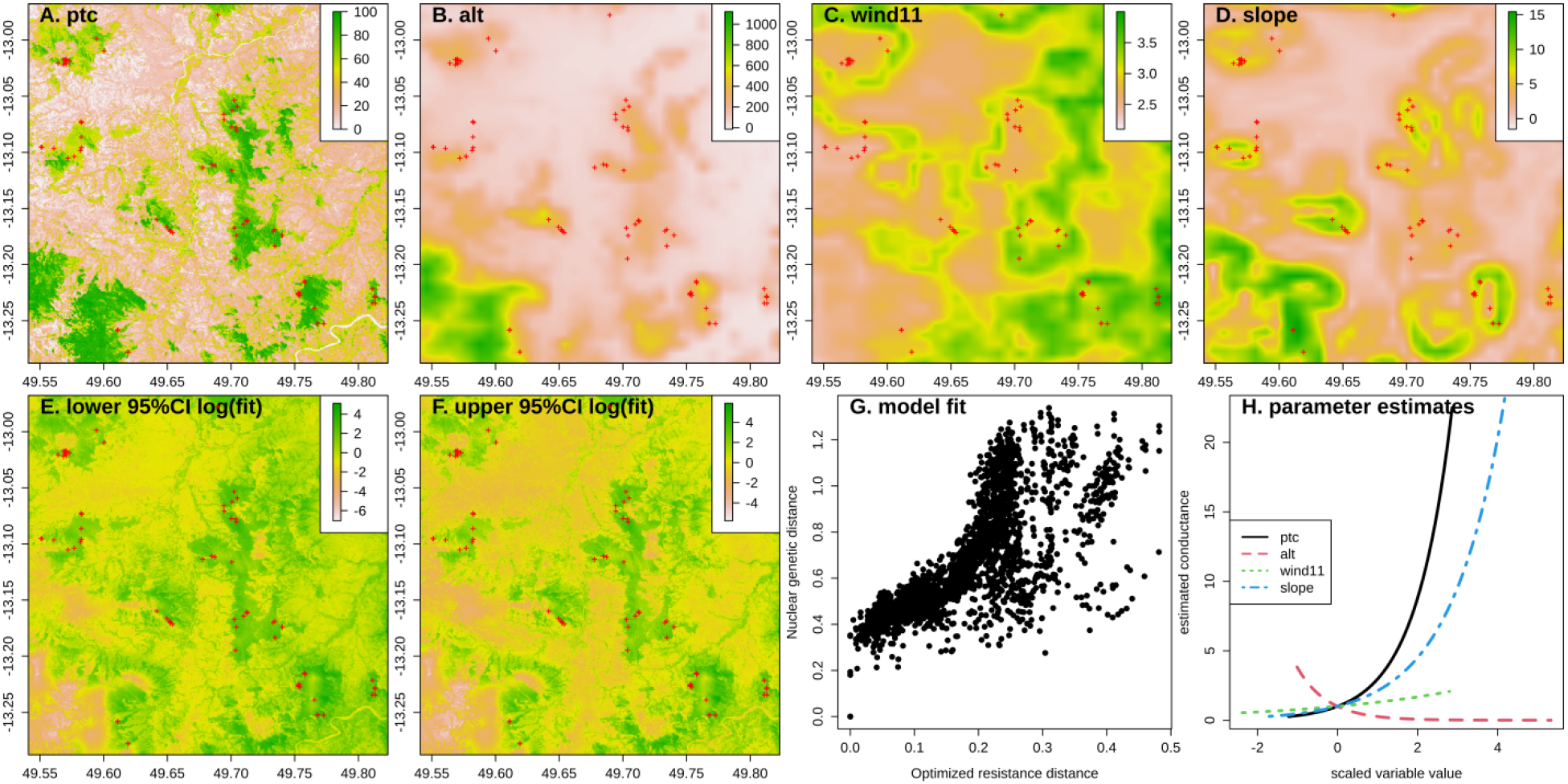
Landscape contribution to nuclear gene flow in *Noronhia spinifolia*. The figure shows the four landscape variables (**A-D**) likely contributing to *N. spinifolia* nuclear gene flow (pollen and seeds), the confidence interval of their fitted composite conductance (**E-F**), the model fit relationship between the optimized resistance distance and the nuclear genetic (**G**), and the relative effects of the four retained variables (**H**). It illustrates a major conducting effect of the current forest cover (ptc: percent of tree cover) on the connectivity of *N. spinifolia*, and it further shows a positive effect of the wind speed in November [wind11] and a composite effect of the topography (positive effect of slope and negative effect of altitude [alt]).

## DISCUSSION

With a comprehensive and extensive sampling of *Noronhia spinifolia* in its core distribution area, and leveraging the rare combination of nuclear and mitochondrial RADseq data with cpDNA microsatellites, this study allowed us to reveal a strong effect of forest cover on gene flow in a patchy habitat in northern Madagascar. We not only report a surprisingly high organellar genetic diversity unevenly distributed in space, but also found that GL-based approaches were able to recover much more information than SNP-calling approaches in our model species. Moreover, the iterative parameter space exploration and the optimization of resistance surface allowed identifying outstanding landscape variables with a strong effect on the connectivity of *N. spinifolia*. Finally, we show that recent forest cover better explains the genetic structure of *N. spinifolia* than more ancient ones.

### *Noronhia spinifolia*, a highly diverse Malagasy microendemic

Our analyses first exhibit unexpectedly high organellar genetic diversity *(h_cp_* = 0.99; 55 chlorotypes for 72 individuals, and *h_mt_* = 0.85; 15 mitotypes) in a microendemic Malagasy tree species. Such cpDNA diversity is tremendously higher than that of another microendemic congener of the High Plateau (*N. lowryi*) when using the same 15 cpSSR loci [6 haplotypes in 77 individuals; *h_cp_* = 0.58 (Salmona *et al*., 2020)]. More surprisingly, more cpDNA haplotypes and diversity were revealed in 72 *N. spinifolia* individuals than in 1263 wild olive trees (*Olea europaea)* from the whole Mediterranean basin [47 chlorotypes; *h_cp_* = 0.35 (Besnard et al., 2013)] and thus across very different geographic scales (LM region = 2,500 km2 *vs* Mediterranean basin = ~2.5 Million km2) and despite the use of more polymorphic cpSSRs (n = 35) in the Mediterranean olive. Similarly, the *N. spinifolia* mtDNA diversity is also higher than in the Mediterranean olive [4 mitotypes; *h_mt_* = 0.58; (Besnard et al., 2002)], although comparable diversity levels have been revealed in other plant groups exhibiting large mitogenomes with high mutation rates as *Silene vulgaris* (Moench) Garcke in Central Europe [30 mitotypes; *h_mt_* = 0.94; (Štorchová & Olson, 2004)]. Secondly, the moderately high levels of nuclear genomic diversity (*H*_E_ = 4.53-6.52 × 10^-3^) are within the range of that estimated in poplar, pedunculate oak and Norway spruce populations across distribution ranges several order of magnitude larger (Chen et al., 2019; Ma et al., 2018; Plomion et al., 2018). Nonetheless, our unexpectedly high species genetic diversity is also comparable to that of other range-restricted microendemics (e.g. Eliades *et al*., 2011; Lanes *et al*., 2018).

Although high standing genetic diversity is generally common in plants, including forest trees as our focal species, the relative importance of the multiple mechanisms generating and maintaining this diversity are still debated (Isabel et al., 2020; Petit & Hampe, 2006; Scotti et al., 2016). In *N. spinifolia*, several non-exclusive evolutionary mechanisms may explain such a high intraspecific genetic diversity. Firstly, it suggests that long-term maintenance of a large effective population size precluded significant genetic drift. Persistent connectivity between forest patches may have been key in this process, particularly during climatic fluctuations of the Late Quaternary that may have contributed to fragmenting habitat, as suggested for other species of the LM region (Salmona, Heller, Quéméré, et al., 2017; Sgarlata et al., 2019). Secondly, the genus *Noronhia* has extremely diversified in northern Madagascar (Hong-Wa, 2016), and about 30 taxa (including already-described species, new species to be described, potential hybrids, and cryptic species) have been recently recorded and sampled in the LM region (JS & GB, unpublished data). What caused such diversification remains unknown. But the co-occurrence of closely related taxa may offer some opportunities for hybridization events, which could have contributed to the increased genetic diversity in *N. spinifolia*. However, the cpSSR characterization of four sympatric/parapatric LM *Noronhia* (i.e. *N. candicans* H.Perrier, *N. clarinerva, N. crassinodis* H.Perrier and *N. intermedia* Hong-Wa; > 200 individuals), closely related to *N. spinifolia* (according to cpDNA and nDNA data; Salmona *et al*., 2020), shows that these species have no shared chlorotype with our study model (GB, unpubl. data), thus suggesting that maternal introgression events to *N. spinifolia*, if any, may not be recent. Lastly, high mutation rate may also contribute to the high genetic diversity in *N. spinifolia*. An obvious acceleration of the mitogenome evolutionary rate has been recently documented in the closely related species *N. candicans, N. clarinerva, N. intermedia* and *N. spinifolia*, with a high number of di- or trinucleotide mutations possibly reflecting frequent mtDNA recombination in this clade (Van de Paer, 2017), as also suggested by a lack of LD between some SNPs (Fig. S2). While accelerated mutation rate was missing on the plastome (Salmona et al., 2020), we are still lacking any evidence for the nuclear genome. Such accelerated evolutionary rate could result from relatively frequent and recurrent hybridization events in this group, promoting genomic instability (Fontdevila, 2005; Payseur & Rieseberg, 2016). Moreover, the strong linear relationship between geographic and genetic distance could preclude cryptic radiation (Pillon et al., 2014) and microgeographic adaptation (Scotti et al., 2016) as major drivers of the observed diversity. In conclusion, the surprisingly high genetic diversity calls for the identification of the evolutionary, ecological and/or molecular mechanisms underlying this peculiar pattern.

### Landscape effects on the genetic diversity of *Noronhia spinifolia*

#### A strong continuous spatial structure

Beyond revealing surprisingly high levels of genetic diversity, our results also show complementary signals of a strong continuous structure in space (PCA, clustering and IBD), from both organelles and the nucleus, in contrast to generally expected incongruent patterns among genomes (Bianconi et al., 2020; Olofsson et al., 2019). While the northwest-southeast differentiation cline represented as much as ~15% of the variance of the PCA on nuclear data, the geographic Euclidean distance alone explained up to ~55% of the nuclear genetic variance using IBD tests. This strong pattern of nuclear genetic structure sharply contrasts with the absence of nuclear spatial structure in the savanna olive tree, *N. lowryi* (Salmona et al., 2020). However, reported IBD patterns in trees show a wide range from low values in *Dalbergia monticola* Bosser & R.Rabev. across eastern Madagascar humid forests [*R*^2^ = 0.18; (Andrianoelina et al., 2009)], or *Coffea mauritiana* Lam. in the Réunion Island [*R*^2^ = 0.21; (Garot et al., 2019)], to high values in *Swietenia macrophylla* King in Central America [*R*^2^ = 0.62; (McRae & Beier, 2007)]. Unexpectedly, this genetic structure was here extremely well explained by the vegetation cover (percent tree cover; mtDNA *R*^2^ = 0.38; nDNA *R*^2^ = 0.90), and to a lower extant by the topography (elevation and slope) and by the wind speed in November. Although strong landscape effects were also found in *S. macrophylla* (McRae & Beier, 2007), we report a unique evidence of a strong habitat effect explained predominantly by one landscape variable.

#### On seed-mediated gene flow: the organellar DNA testimony

Although organellar IBR patterns (Figs S15, S18-20, S24) suggest that seed-mediated gene flow is driven by forest cover, the recovered pattern was of lower intensity than for pollen-mediated gene flow (nDNA). Despite watershed networks being candidates for hydrochory, only slope (barochory) and altitude showed noticeable effect on seed dispersal. Similarly, the overall structures of organellar haplotype networks (Figs 2, S12-S13) are coherent with the geographic repartition of forests, and in line with the effect of the forest cover. These prevailing effects of forest cover, and the limited haplotype sharing between forests, suggest that seed dispersal may be relatively limited, primarily driven by slope (barochory) and performed by forest-dwelling animals (zoochory), especially those with limited and/or rare trans-forest movements, such as lemurs, rodents and territorial birds (Aleixo-Pais et al., 2019; Quéméré et al., 2010; Rakotoarisoa et al., 2013a; Sgarlata et al., 2018). Furthermore, the limited haplotype sharing between forests suggest that *N. spinifolia’*s organellar diversity is the result of mutation-drift rather than that of dispersal-drift processes (Cruzan & Hendrickson, 2020). However, the networks also show multiple potential fluxes among forests, hence recalling the network complementarity to the IBR approach. Several non-exclusive interpretations can be invoked for explaining these patterns: (*i*) relevant landscape variables are not included or of low resolution (e.g. watershed networks and climatic variables); (*ii*) the cpDNA and mtDNA diversities are confounded by homoplasy, recombination, strong drift, long-term phylogenetic or demographic history; (*iii*) seed dispersal also results from infrequent seed ingestion by wide-ranging birds (or other vertebrates); and (*iv*) the sampling possibly limited resolution at the seed dispersal scale.

#### A deep forest cover effect on gene flow

Unlike organellar DNAs, nDNA diversity is deeply explained by the LM region forest cover (Fig. 4). While this partially confirms the effect of forest cover, elevation and slope on seed dispersal, since nDNA diversity is influenced by both seed and pollen movement, wind-mediated pollen dispersal is also supported. Indeed, November wind speed, which fits the flowering temporal window of *N. spinifolia* (Hong-wa, 2016), shows a subtle but significant effect (Fig. 4) suggesting that pollen dispersal or pollinator movements could be influenced by the wind regime. Combined, the strong forest cover effect, the slope, elevation and wind effects suggest that pollen dispersal could be mediated by forest-dwelling organisms with movements limited by non-forest environments and high elevation, but facilitated by slope and November wind speed. Insect-mediated pollen dispersal in *N. spinifolia* is also strongly suggested by its flower morphology and color (Hong-Wa, 2016). However, the currently limited knowledge of the Malagasy entomofauna and plant-pollinator networks prevents us from clearly identifying this species’ forest-dwelling pollinators. Furthermore, our analyses are limited by (*i*) the lack of high-resolution wind directionality data, and (*ii*) their lack of integration (Fernández-López & Schliep, 2018; Kling & Ackerly, 2021) with composite landscape surface optimization approaches (Peterman, 2018; Peterman & Pope, 2021).

### The antiquity of habitat mosaic in northern Madagascar

Our results further support a long-standing habitat mosaic in the LM region. First, the better fit of the recent forest cover (~2000s), compared to older vegetation cover (~1953, ~1973), suggests that the small forest changes that have occurred through this period (Quéméré et al., 2012) are unable to explain the genetic diversity of *N. spinifolia*. These mild landscape changes, mostly located at the forest patches edges (Quéméré et al., 2012), contrast with the high deforestation rates observed throughout Madagascar since the 1950s. They might be explained by the low human settlement density (Green & Sussman, 1990; Schüßler et al., 2020) and the dominant dry forest type (Hansen et al., 2013; Vieilledent et al., 2018) of the LM region. Under such high recent deforestation rates, a better fit of the recent forest cover layer would be very unlikely, even considering that its better resolution could positively bias its fit. Second, because we mostly genotyped fully-grown mature trees, and since the generation time of *Noronhia* is potentially long [>20-50 years; (Salmona et al., 2020)], the genetic diversity is expected to reflect ancient forest cover. The time lag for a particular landscape feature to imprint its effects in the genetic diversity of a species, has been little studied (Landguth et al., 2010; Mona et al., 2014). However, in *N. spinifolia*, based on the strength of the signal, the high level of diversity and of gene-flow, the re-shuffling of allele frequencies after fragmentation can be relatively long (tens to hundreds of generations), before harboring the signature of the new geographical pattern. The period with data on forest cover (1953-present) represents less than five generations, a too short period to erase the signal of previous population structure (or lack thereof). This suggests that the landscape changes leading to the current forest cover long predates the most ancient available layer (1953). The strong genetic correlation with the recent forest cover is, therefore, sound evidence that the landscape of the LM region was relatively stable at least for the last century (i.e. when most of Madagascar’s deforestation occurred), and possibly the last millennium. This result concurs with those of recent studies (Quéméré et al., 2012; Salmona et al., 2020) supporting a relative antiquity of habitat mosaic in northern Madagascar. Furthermore, both the high diversity of *N. spinifolia*, and its predominant distribution in low-elevation dry forest suggests that this habitat type may have been spatially, topographically, and temporally extensive in northern Madagascar, albeit frequently fragmented, as seemingly evidenced by a rare and likely relictual occurrence of the species in contemporary high-elevation humid forest (e.g. Binara) and similarly peculiar presence further north (e.g. Montagne des Français). To assess forest-cover changes over a larger time frame (e.g. the last ten or so millennia), inferences of *N. spinifolia”*s demography over time would be relevant (Beichman et al., 2018; Salmona, Heller, Lascoux, et al., 2017). Coupling these inferences, with that of short-generation grassland organisms, would also help clarifying the dynamics of fire-prone non-forest environments, through the succession of environmental changes that occurred during last millennia, namely the last-glacial-maximum, early human’s colonization, the mid-Holocene transition, and the 1-Kya expansion of agro-pastoralism.

### Further prospects and conservation implications

The power of coupling genomic data to landscape genetics allowed not only identifying major landscape components influencing effective dispersal, but also their respective effects on seed and pollen dispersal. This surprising result warrants further investigation using higher resolution landscape and environmental layers, not used, or not available to our study. In particular, it would benefit from the use of forest type, soil type, land use, and climate data of better resolution. In addition, the wind effect has been inferred without considering its directionality. Recent landscape genetics analytical advances allowing directionality integration (Fernández-López & Schliep, 2018; Kling & Ackerly, 2021) are still limited by the lack of high-resolution directional data (e.g. wind), and of integration with composite landscape surface optimization approaches (Peterman, 2018; Peterman & Pope, 2021). These analyses may also benefit from the use of genetic distances reflecting the directional changes in diversity among pairs of samples. Recent methodological development contrasting with previous evaluations of the MLPE (Peterman et al., 2019; Shirk et al., 2018; Winiarski et al., 2020) showed that the Generalized Wishart has higher accuracy in identifying 2-variables composite landscape surface (Peterman & Pope, 2021). Although this is beyond the scope of our study, cross-validation approaches could assess this new optimization model, the effect of seemingly clustered sampling of *N. spinifolia*, or that of modifying the conductance function (Peterman, 2018). While our study clearly identifies that seed and pollen are dispersed by forest-dwelling organisms, it neither identifies these organisms nor does it clearly show that seed and pollen do still effectively disperse among forests. These questions could be tackled (i) by inferring pedigree data from high density population sampling, coupled with sampling of young trees and seedlings, (ii) using field survey of potential dispersers during flowering and fructification (e.g. camera tracking), and/or (iii) using metabarcoding approaches to assess the interaction network within the LM forests. While our study confirms the biological importance of the LM region, which is known for its species richness and endemism across taxa (Goodman & Wilmé, 2006; Rakotondravony, 2006, 2009; Sgarlata et al., 2019), and more specifically for the genus *Noronhia* (Hong-Wa, 2016), our results also have several implications for biodiversity conservation in the region:

- First, they underscore the conservation value of the often-overlooked intraspecific genetic diversity, which is unexpectedly high in *N. spinifolia*.
- Second, this study highlights the importance of riparian forests of the LM region for their major role both as corridors connecting forest patches, which is supported by the fact that genetic diversity in *N. spinifolia* is explained by forest cover rather than geographical distance, and as vectors promoting the roles of vertebrates and insects on seed and pollen dispersal. Therefore, actively maintaining, protecting, and reforesting riparian and corridor forests, which are likely pivotal for the functional connectivity of *N. spinifolia* but also most native and endemic species of the LM region (Aleixo-Pais et al., 2019; Rakotoarisoa et al., 2013a; Sgarlata et al., 2018), remain critical conservation actions.
- Third, our study identifies the Binara forest as unique among the major forests of the LM region and in urgent need of deeper conservation focus. Indeed, our extensive forest survey allowed us to find and collect just a few *N. spinifolia* samples in this forest, where they were found only at unexpectedly higher altitude and wetter habitat (Fig. S1). Similarly, several other Malagasy olive species that are mostly distributed in dry forests [e.g. *N. ankaranensis* (H.Perrier) Hong-Wa, *N. candicans*, *N. christenseniana* Hong-Wa and *N. oblanceolata* H.Perrier; GB and JS unpublished data], were also found to occur only at higher altitude in the mountain evergreen forests of this region (e.g. Binara and Antsahabe). Altogether, this pattern, though unclear, echoes the peculiarities of these forests that likely acted as refugia for numerous taxa during drier periods (Goodman & Wilmé, 2006; Rakotoarisoa et al., 2013b; Raxworthy & Nussbaum, 1995; Sgarlata et al., 2019).

### Data accessibility

Raw RADseq data and RADseq mtDNA alignments have been deposited at the Short Read Archive (SRA) NCBI database under the reference PRJNA632767. Organellar microsatellite genotypes and mtRAD variants are available in Tables S9 and S10, respectively. All additional data, scripts and materials are available to readers at https://doi.org/10.5281/zenodo.7193354.

### Benefit sharing

Benefits Generated. Research collaborations were developed and strengthened with scientists from Madagascar, the country of origin of the genetic samples. One Malagasy student (AER) was trained during this work. All academic collaborators are included as co-authors. Three guides from the Loky-Manambato protected area protected area guides and cook association were trained to botanical work, as well as several additional guides in remote communities. The research results have been and will be repeatedly shared with the concerned communities, the managers of the protected area, and the broader scientific community. The research also addresses a priority concern, in this case the conservation of the focal organism, *N. spinifolia*,and by extent its habitats, the Loky-Manambato region and Northern Madagascar. More broadly, our group is committed to international scientific partnerships, as well as institutional capacity building.

## Acknowledgments

We thank the Direction Générale du Ministère de l’Environnement et des Forêts de Madagascar, Madagascar’s Ad Hoc Committee for Fauna and Flora, and Organizational Committee for Environmental Research (CAFF/CORE) for permission to perform this study (permits number: [49/17] & [127/18]/MEEF/SG/DGF/DSAP/SCB.Re) and for their support. We thank the local communities of Daraina and of the LM region for their warm reception and support. We thank E. Rasolondraibe and the many local guides and cooks for sharing their incomparable expertise and help in the field, *misaotra anareo jiaby*. We thank U. Suescun, C. Verbeke and M.A. Naranjo Arcos for lab assistance, L. Chikhi for comments on an earlier version of the draft. This work was mostly funded through an ERA-NET BiodivERsA project: INFRAGECO (Inference, Fragmentation, Genomics, and Conservation, ANR-16-EBI3-0014). We also thank the LABEX TULIP (ANR-10-LABX-0041) and CEBA (ANR-10-LABX-25-01), and the LIA BEEG-B (Laboratoire International Associé – Bioinformatics, Ecology, Evolution, Genomics and Behaviour, CNRS). We are grateful to the Get-Plage sequencing, Genotoul bioinformatics (Bioinfo Genotoul) and GenOuest bioinformatics platforms for sequencing services and providing computing resources. The fourth preprint version of this manuscript has been peer-reviewed and recommended by Peer Community In Evolutionary Biology (https://doi.org/10.24072/pci. evolbiol.100136).

## Author Contribution

JS and GB designed the experiment. JS, AER, BLP, JR, CHW and GB were pivotal to field material collection and herbarium composition. JS, SM, and GB generated the genetic data. JS conducted bioinformatics and population genetic analyses. JS and AD conducted IBR analyses. JS and GB drafted a first version of the manuscript with significant input from CHW. All coauthors agreed with the last version of the manuscript.

