## Supporting Information for "How ancient forest fragmentation and riparian connectivity generate high levels of genetic diversity in a microendemic Malagasy tree"

#### Table of content:

Supporting methods 3

- Method S1: DNA extraction 3
- Method S2: Chloroplast microsatellites 3
- Method S3: Organellar markers processing 3
- Method S4: RAD sequencing 4
- Method S5: Screening the organellar genomes for RADseq loci 4
- Method S6: *De-novo* assembly of the nuclear loci catalog and ploidy assessment. 4
- Method S7: SNP calling & Genotype likelihood 5
- Method S8: Clustering analyses 5
- Method S9: Isolation by distance 5
- Method S10: Landscape variables, cost and resolution 6

Supporting results 7

- Result S1: Organellar DNA genotyping 7
- Result S2: Catalog construction and genotypes data 7
- Result S3: Isolation by distance 7

Supporting tables 8

- Table S1: Samples and genetic data used in this study 8
- Table S2: Organellar optimized composite landscape conductance models 8
- Table S3: Nuclear optimized composite landscape conductance models 8
- Table S4: Characteristics of the organellar DNA microsatellites 9
- Table S5: Characteristics of the mtDNA variants obtained from RADseq data (mtRAD) 10
- Table S6: Organellar $F_{ST}$ among sampling sites 11
- Table S7: Organellar Nei and Edward's *D* among sampling sites 12
- Table S8: Nuclear genetic differentiation among *Noronhia spinifolia* sampling sites 13

Table S9: Organellar microsatellite genotypes 14
Table S10: Mitochondrial RADseq genotypes 14
Supporting figures 15
Figure S1: Altitudinal distribution range of *Noronhia spinifolia* in the Loky-Manambato region 15
Figure S2: Linkage disequilibrium in organellar DNA data 16
Figure S3: Number of nuclear genetic clusters best explaining the data when using NgsAdmix 17
Figure S4: Number of nuclear genetic clusters best explaining the data when using Admixture 18
Figure S5: Genetic structure in *Noronhia spinifolia* 19
Figure S6: ngsAdmix ancestry proportion estimates for $K$ = 2 to 10 20
Figure S7: Admixture ancestry proportion estimates for $K$ = 2 to 10 21
Figure S8: Geographic scale influence on isolation by distance (IBD) 22
Figure S9: *Noronhia spinifolia*'s genetic diversity 23
Figure S10: Spatial distribution of nuclear genetic diversity in *Noronhia spinifolia* 24
Figure S11: Altitude effect on *Noronhia spinifolia*'s genetic diversity 25
Figure S12: *Noronhia spinifolia* mtDNA haplotype network 26
Figure S13: *Noronhia spinifolia* chlorotype network 27
Figure S14: Forest based organellar isolation by distance in *Noronhia spinifolia* 28
Figure S15: Isolation by distance in *Noronhia spinifolia* 29
Figure S16: Mantel correlogram of spatial correlation 30
Figure S17: Principal component analysis of nuclear genomic data of *Noronhia spinifolia* 31
Figure S18: Univariate variable selection for chloroplast data 32
Figure S19: Univariate variable selection for mitochondrial data 33
Figure S20: Univariate variable selection for organellar data 34
Figure S21: Univariate variable selection for nuclear data 35
Figure S22: Univariate optimized association with genetic distances. 36
Figure S23: Univariate monthly wind speed association with nuclear genetic distances. 37
Figure S24: Landscape contribution to organellar gene flow in *Noronhia spinifolia*. 38
References 39

### Supporting methods

#### Method S1: DNA extraction

Out of 220 collected *N. spinifolia* trees, we extracted DNA for 137 samples, selected to maximize geographic and altitudinal representation and prioritizing mature, tall and large trees. We ground 0.5 cm² dried leaf pieces in a 2-mL tube with three iron beads with a TissueLyser (Qiagen). We extracted total genomic DNA using the BioSprint 15 DNA Plant Kit© (Qiagen) according to the manufacturer's protocol and finally eluted DNA in 200 µL AE buffer. We quantified nucleic acids using the Nanodrop spectrophotometer (ND-1000, Thermo Fisher) and double stranded DNA (dsDNA) using a Qubit 2.0 Fluorometer (Invitrogen). For an optimal genetic characterization, a dsDNA concentration superior to 4 ng/µL and a 260/280 absorbance ratio superior to 1.75 were required. Finally, by optimizing both geographic and altitudinal ranges, 72 samples were selected for RAD sequencing (Fig. 1).

#### Method S2: Chloroplast microsatellites

Chloroplast microsatellites (cpSSR) were amplified in a multiplex of three or four loci using the method described by Schuelke (2000). We targeted cpSSRs because such markers are expected to be the most variable parts of the plastome and may inform on relatively recent demographic processes (e.g. Besnard et al., 2011, 2013). A total of 15 cpSSRs were used (Table S1). These loci were originally developed on the Olive tree (*Olea europaea*; Besnard et al., 2011). We selected 15 loci based on full plastome sequences of several *Noronhia* species, including one *N. spinifolia* (Olofsson et al., 2019; Salmona et al., 2020). We first checked that primer sites were conserved between the *Olea* and *Noronhia* genera, and then considered loci with a poly-T stretch containing at least ten repeats. We amplified the 15 cpSSR loci in four multiplex reactions, using four distinct fluorochromes (Table S2). All loci were genotyped together, within one multiplexed run, on an ABI Prism 3730 DNA Analyzer. Note that one primer pair (cpSSR locus n°32, Table S2) amplified an additional polymorphic paralogous SSR locus from the mitogenome (mtSSR; see below).

#### Method S3: Organellar markers processing

For each individual, allele size at each cpSSR locus was scored. A haplotype (or chlorotype) was then defined by combining alleles at the 15 loci. Multi-state microsatellites were then treated as ordered alleles and coded by the number of repeated motifs for each allele [e.g., number of T or A (Besnard et al., 2011)] considering as a reference the plastome of individual GB124-2017 (MT081057). One variable mtSSR [due to an *mtpt* region homologous to cpSSR locus 32 (Van de Paer et al., 2018)] was also amplified. It was independently scored and compared to other mitochondrial loci (mtRAD; see below).

Organellar genomes are supposedly inherited from the mother in the olive tribe (Van de Paer et al., 2018), but some exceptions have been reported in the core Ligustrinae (Liu et al., 2004). As we cannot directly verify this assumption on progenies, we here investigated the linkage among organellar markers that may unravel the presence of potential events of recombination. The Agapow & Burt $r_d$ linkage disequilibrium (LD; Agapow & Burt, 2001) using the package *poppr* (Kamvar et al., 2014, 2015) was estimated among cpSSR, mtDNA loci and among all organellar markers.

To assess the effect of combining organellar data, we tested cpSSR / mtRAD genetic distances ratio ranging from 1 to 20 and evaluated their fit to the IBD model. We retained the best fitting maximum pairwise geographic Euclidean distance (S) between samples by 2,000-m increments. At this S distance, we retained the best fitting cpSSR / mtRAD ratio (see Method S10 and Fig. S25 for further details).

#### Method S4: RAD sequencing

Restriction site-Associated DNA sequencing (RADseq) consists in sequencing regions neighboring restriction sites, to obtain homologous sequences across many individuals, spread across the genome, with a decent coverage and at a reasonable cost (Andrews et al., 2016; Baird et al., 2008). We prepared RADseq libraries using 100-200 ng of genomic DNA from each sample and one negative control made up of water following the protocol described in Etter et al. (2012) and by the Genomic Resources Development Consortium (2015). Briefly, samples were digested with *Sbf*I (New England Biolabs) before P1 adapter ligation. After adapter ligation, we pooled 10 µL of 18 samples in four sub-libraries, before shearing them in a Covaris M220 for 45 sec to obtain fragments with an average size of 500 bp. Following end repair, we conducted size selection using AMPure XP beads (Agencourt). After 3'adenylation, we ligated the P2 adapter. The libraries were finally enriched with 10 cycles of PCR. Libraries quality was verified using Fragment Analyzer® (Advanced Analytical) and quantified using qPCR (QuantStudio6®, Applied Biosystems). We made equimolar pooling with the sub-libraries and sequenced them on an Illumina HiSeq-3000 (150-pb paired-end reads; 72 individuals / lane) at the GetPlage Sequencing platform facility (Toulouse, France). Raw reads were then demultiplexed in Stacks v2.0b (Rochette et al., 2019) using default parameters and quality-filtered using Trimmomatic (Bolger et al., 2014) with the following parameters: Leading: 3, Trailing: 3, Slidingwindow: 4:15, Minlen: 60.

#### Method S5: Screening the organellar genomes for RADseq loci

An initial *in silico* text mining survey of *Sbf*I restriction sites (grep function in bash) in *N. lowryi* plastome (Olofsson et al., 2019; Salmona et al., 2020) and *N. clarinerva* mitogenome [MW202230; assembled following (Van de Paer et al., 2018)] revealed no *Sbf*I site in the chloroplast genome, while ten sites were found in the mitogenome. We thus first recovered mtDNA information from RAD cleaned sequence bwa-mem alignment (Li, 2013) to the *N. clarinerva* mitogenome. The ten *Sbf*I RAD loci identified *in silico* were recovered at high coverage (average individual forward RAD sequence depth = 22,000×, Table S1, S3) in alignment files. Haplotypes were called using ANGSD v0.92 (Korneliussen et al., 2014; Nielsen et al., 2012) based on their highest effective base depth (Wang et al., 2013), limiting our analyses to region with at least 50× coverage and to 700-bp windows on each side of the ten *Sbf*I cutting sites. One large indel on the $10^{th}$ loci limited us to only using its last 900 bp. In contrast, no *Sbf*I cpDNA RAD locus could be recovered in the *Noronhia* plastome, confirming our *in silico* initial analyses.

#### Method S6: *De-novo* assembly of the nuclear loci catalog and ploidy assessment.

A catalog of tags (loci) was *de-novo* optimized, from a limited number of individuals (eight), spread across all sampled forests, by iterating around and selecting the core parameters for Stacks (*m*, *M* and *N*) to maximize the amount of available biological information (Paris et al., 2017). We allowed the parameters *m* – the minimum number of reads required to build a stack – to vary between 2 and 8, *M* – the maximum number of differences between stacks of an individual allowed when building a locus – from 2 and 8 and *N* – the maximum number of differences between loci of multiple individuals allowed when building a loci – from 2 to 8. The final catalog built with Stacks (selected parameters: m = 4, M = 5, N = 8) from a larger set of individuals (21) spread across all sampled forests, was decontaminated using DeconSeq (Schmieder & Edwards, 2011) by assessing sequence similarity to Bacteria, Archaea, Virus, Salmonella, and Human databases of potential contaminants, in comparison with the Rice database (closest relative in Deconseq, i.e. non-contaminant). To minimize allelic dropout bias from loci under-merging, orthologous loci across the catalog were identified using MUMmer (Kurtz et al., 2004) and removed if above a threshold of 88% of homology. We also removed sequences that may not originate from the nuclear genome, i.e. loci with high homology to the *Noronhia clarinerva* mitogenome (MW202230) as well as plastomes of *N. lowryi* (NC_036984) using MUMmer as mentioned above.

Ploidy was first inspected using minor allele frequency plots. Ploidy was further statistically confirmed using nQuire (Weiß et al., 2018) with default parameters, a minimum depth of 10, a minimum mapping quality of 60, and using all implemented statistical tests.

##### Method S7: SNP calling & Genotype likelihood

We used two fundamentally distinct genotyping approaches to ensure the robustness of our results: single nucleotide polymorphism (SNPs) called in Stacks, and genotype likelihoods (GLs) estimated with ANGSD (Methods S7). GLs retain information about uncertainty in base calls, which alleviates some issues associated with RADseq data such as unevenness in sequencing depth and allele drop-outs (Heller et al., 2021; Pedersen et al., 2018; Warmuth & Ellegren, 2019).

**SNP**: We called genotypes with the *populations* module of Stacks (Rochette et al., 2019) using the final catalog generated in Method S6, and discarding loci present in less than 20% of the 72 individuals. Using VCFtools v0.1.15 (Danecek et al., 2011), we filtered-out SNPs with a depth <10× and >250×, a mean depth across all individuals <5×, a genotype quality <20, minor allele frequencies <0.05, and more than 90% missing data. We also kept only biallelic SNPs. Subsequently, using PLINK-0.1.15 (Purcell et al., 2007), we selected one SNP per locus to prevent linkage disequilibrium from introducing a bias in downstream analyses.

**GL**: Cleaned sequences were aligned using bwa-mem (Li, 2013) to the *de-novo* catalog of nuclear, decontaminated, non-orthologous loci. We estimated loci depth as the F1 read starting position (*Sbf*1 cutting site) using the "samtools-view" command (Li et al., 2009) in R (R CoreTeam, 2014). We then estimated genotype likelihoods ("-GL 1") using only loci with (1) a total depth of at least twice the number of individuals ("setMinDepth"), (2) a total depth of at most ("-setMaxDepth") the sum of the 0.95 quantile of the depth distribution of considered individuals, and (3) an individual depth of at most ("-setMaxDepthInd") the maximum 0.95 quantile of considered individuals. This strategy allows discarding loci present only in a very low number of individuals, or over-covered loci that are likely to be repeated or paralog regions (Heller et al., 2021; Pedersen et al., 2018). In addition, we considered only bases with a minimum quality ("-minQ") of 20, reads mapping uniquely ("-uniqueOnly") with a minimum quality ("-minMapQ") of 30 and associated with their pair ("-only_proper_pair"; for PE only). Finally, we only kept biallelic variants ("-skipTriallelic") with a probability ("-SNP_pval") below 1e-6 and a minor allele frequency ("-minMaf"; MAF) of at least 0.05.

##### Method S8: Clustering analyses

To explore the genetic structure of our study system, we performed naive clustering analyses based on ANGSD genotype likelihoods using NgsAdmix v32 (Skotte et al., 2013) and on Stacks called genotypes using ADMIXTURE v1.3.0 (Alexander et al., 2009). Admixture proportions were inferred with the number of clusters ($K$) ranging from two to the number of hypothesized populations (here forest patches) plus two (i.e., 2-10). NgsAdmix was run for 20 iterations per $K$ value and a stringent tolerance of $10^{-6}$ for convergence. We compared $K$ values using the $\Delta K$ procedure (Evanno et al., 2005). ADMIXTURE was run using the default optimization method and the default termination criterion that stops the analysis when the log-likelihood increases by less than $10^{-4}$ between iterations. $K$ values were compared using the cross-validation method ("--cv" flag in ADMIXTURE) and by comparing likelihoods across runs.

##### Method S9: Isolation by distance

We investigated patterns of isolation by distance (IBD) to assess how the geographic distance alone explains the genetic diversity (Slatkin, 1993; Wright, 1943). To that respect, we used Mantel tests (Mantel, 1967) between individual geographic and genetic distances, with 9999 permutations, using the *ade4* R package (Chessel et al., 2004). Since IBD may be limited to a certain scale (e.g. Keller &

Holderegger, 2013; Van Strien et al., 2015), we compared subsets of pairwise data defined by a maximum geographic distance (S) between samples (Cayuela et al., 2019). S ranges from 9,000 m [the estimated distance at which no *N. spinifolia* individual is excluded from a neighboring graph (Jombart, 2008)] to 35,000 m (the maximum Euclidean distance between two individuals in our study), by 2,000 m increments. For each subset, we ran the IBD test and plotted the model fit ($R^2$) against S to assess which spatial scale optimizes the amount of variance explained (Cayuela et al., 2019).

##### Method S10: Landscape variables, cost and resolution

As *N. spinifolia* was recently described and occurs in a remote area (Hong-Wa, 2016), we had little prior knowledge on the landscape variables that may affect pollen and seed dispersals. We therefore assessed the effect of most available landscape variables for the study region. We considered roads, trails, rivers, streams, slope (https://www.diva-gis.org/gdata), and wind speed (Fick & Hijmans, 2017; http://worldclim.org/version2). The slope was estimated from the elevation data using the *raster* R package (Hijmans et al., 2015). We further assessed the effects of the vegetation cover. Considering that the genetic diversity of the old trees may be better explained by past forest cover, we used categorical forest cover data from 1953, 1973, 2000s (Vieilledent et al., 2018; https://bioscenemada.cirad.fr/maps/), and continuous percent tree cover from 2000s (Hansen et al., 2013; http://earthenginepartners.appspot.com/science-2013-global-forest). The latter was considered continuous (as is) or discrete (only cells with >40% tree cover considered as forest). Altogether, we assessed the effect of five discrete and seven continuous landscape variables (Table 1).

Although strong priors associating a landscape component to a particular cost may be available for well-studied species (e.g. Dellicour et al., 2019; Quéméré et al., 2010), landscape variables and their associated cost are often chosen almost arbitrarily when little or no data are available (Beier et al., 2008, 2011). To identify the variable-cost associations that matter for our study system, we iteratively tested 14 conductance-resistance values [1:20, 1:15, 1:10, 1:8, 1:5, 1:4, 1:2, 2, 4, 5, 8, 10, 15, 20]. Each of these 14 values is assigned to the landscape feature (cost of the variable). A value of “1” is assigned to the rest of the surface (cost of the non-variable). A resistance surface is created from each combination of two values. Values below and above one represent conductance and resistance, respectively. Similarly, organisms do not necessarily perceive each environmental component at the same resolution (or granularity: Baguette & Van Dyck, 2007; Everson & Boucher, 1998; Laurance et al., 2007; Murcia, 1995). To identify the variable-cost-granularity relevant for *N. spinifolia*, we tested four pixel resolutions [1-6].

### Supporting results

##### Result S1: Organellar DNA genotyping

Out of the 15 chloroplast microsatellites, all except cpSSR-9 showed polymorphism, with a number of alleles per locus ranging from 2 to 8 (Table S2). Their combination allowed us to distinguish 55 chlorotype profiles among 72 trees. The ten mitochondrial RAD loci (mtRAD) identified *in silico* and recovered at very high coverage (Table S1) allowed identifying 17 SNPs. Interestingly, SNPs were not randomly distributed along RAD locus sequences, with two di- and two tri-nucleotide mutations. A variable mtSSR [due to an *mtpt* region homologous to cpSSR-32 (loc. 679 985-680 084; GenBank no MW202230)] was also scored with three length variants (Table S2). The combination of mtRADs and the mtSSR locus leads to the identification of 15 mitotypes among 72 trees (Table S3).
High and significant LD values (>0.4) suggest that the markers are linked and non-recombining. However, low and non-significant LD values are more complex to interpret because these can result from homoplasy, recombination, and/or low diversity at the investigated loci (Mueller, 2004). Both cpDNA and mtDNA gave contrasting signals (Figs S2-S4). The cpSSR markers showed low to moderate LD (Fig. S2). Reduced LD could partly be the consequence of microsatellite-repeat-length homoplasy. Meanwhile, the mtDNA SNP markers showed either complete (among seven loci) or no LD (Fig. S3), the latter indicating putative events of recombination that may result from occasional paternal leaks of mitochondria.

##### Result S2: Catalog construction and genotypes data

From more than 500 million reads (between 1.8 and 36 million reads/ind; average = 7.8 million reads/ind; Table S1), the nuclear catalog parameter space exploration allowed selecting values (m = 4, M = 5, N = 8) that offer a trade-off between the coverage, the number of loci, and the number of SNPs, while limiting the number of paralogs and the presence of contaminants. Briefly, this final catalog combined nuclear loci for which identical reads were found at a minimum coverage of 4×, with a maximum of five differences in same-individual alleles and of eight differences in across-individuals' alleles. The final cleaned catalog was composed of 50,361 nDNA loci, with a F1 mean individual coverage comprised between ~9 and ~68× (Table S1). The final filtered data sets comprised 170,799 SNPs (with ~87% missing data) and 14,580 linkage disequilibrium filtered SNPs (with ~91% missing data) obtained using Stacks and 19,043 genotype likelihood (GL) obtained using ANGSD. The SNP-calling procedure showed low ability to recover the genetic makeup of *N. spinifolia* (when compared to the GL-based procedure; Figs S14, S15, S18), we therefore limited its use to preliminary analyses (ADMIXTURE & genetic distances) and proceeded with the GL-based procedure for downstream analyses.

##### Result S3: Isolation by distance

Organellar and nuclear IBD signals were always maximized when considering the whole sampling distribution (Fig. S13). The geographic Euclidean distances showed low, but highly significant, power at explaining genetic distances among individuals (R2 [cpSSR]: 11.7%; R2 [mtRAD]: 20.7%; and R2 [cpSSR + mtRAD]: 21.3%; Fig. S19). We found a clear IBD signal explaining up to 56.6% of the among-individuals nuclear GL covariance (Fig. S19).

### Supporting tables

##### Table S1: Samples and genetic data used in this study

*Also available in bioRxiv:* https://www.biorxiv.org/content/10.1101/2020.11.25.394544v4
Accession ID: Sample identifier; Forest: local name of the forest; Site: sampled site (several per forest). Latitude and Longitude are given in WGS 1984, degree decimal; RAD index: barcode; Raw read: number of total initial raw reads; Retained reads: number of reads retained by process_radtag after removing reads without the cutting site; De novo it.: individual used for the *de novo* catalog building parameter space exploration; De novo ref.: individual used for the final *de novo* reference building; y: included; na: not included; Mean cov.: average coverage; $H_E$: genetic diversity estimated as the proportion of heterozygous loci from genotype likelihood based on folded site frequency estimated in ANGSD. #: number.

##### Table S2: Organellar optimized composite landscape conductance models

*Also available in bioRxiv:* https://www.biorxiv.org/content/10.1101/2020.11.25.394544v4
Table of organellar optimized composite landscape conductance models assessed in *radish* (Peterman & Pope, 2021). Cov1-2-3-4: first-fourth landscape covariate tested, nb: the order of covariates does not matter. Grain: pixel-size reduction, all model comparison have been conducted with the same maximal grain, conductance: conductance model in *radish*, measurement: measurement model in *radish*, aic:Akaike information criterion, df: degree of freedom, cov1-4_estimate: model posterior estimate associated with the respective covariate, cov1-4_pval: *p-value* associated with the respective covariate estimate, cov1-4_z_Std_Error: model posterior standard error associated with the respective covariate estimate, cov1-4_z_value: model posterior z-value associated with the respective covariate estimate.

##### Table S3: Nuclear optimized composite landscape conductance models

*Also available in bioRxiv:* https://www.biorxiv.org/content/10.1101/2020.11.25.394544v4
Table of nuclear optimized composite landscape conductance models assessed in *radish* (Peterman & Pope, 2021). Cov1-2-3-4: first-fourth landscape covariate tested, nb: the order of covariates does not matter. Grain: pixel-size reduction, all model comparison have been conducted with the same maximal grain, conductance: conductance model in *radish*, measurement: measurement model in *radish*, aic:Akaike information criterion, df: degree of freedom, cov1-4_estimate: model posterior estimate associated with the respective covariate, cov1-4_pval: *p-value* associated with the respective covariate estimate, cov1-4_z_Std_Error: model posterior standard error associated with the respective covariate estimate, cov1-4_z_value: model posterior z-value associated with the respective covariate estimate.

**Table S4: Characteristics of the organellar DNA microsatellites**

Fifteen chloroplast microsatellites loci [cpSSRs (Besnard et al., 2011)] plus one mitochondrial SSR locus (mt-32) were amplified in four multiplex reactions (PCR mx.), each with a given fluorochrome (Flu.). For each fluorochrome, the allele size range (Allele size) of each locus was not overlapping, allowing us to genotype all 16 loci within one multiplexed run, on an ABI Prism 3730 DNA Analyzer.

| Marker ID | Allele size | Flu. | PCR mx. | Number of alleles |
|---|---|---|---|---|
| cp-32 | 106-107 | YAK | 1 | 2 |
| mt-32 | 113-119 | YAK | 1 | 2 |
| cp-9 | 129 | YAK | 1 | 1 |
| cp-22 | 154-157 | YAK | 1 | 4 |
| cp-17 | 177-179 | YAK | 1 | 3 |
| cp-19 | 106-110 | AT565 | 2 | 5 |
| cp-15 | 136-144 | AT565 | 2 | 8 |
| cp-25 | 178-179 | AT565 | 2 | 2 |
| cp-46 | 111-113 | AT550 | 3 | 3 |
| cp-42 | 138-140 | AT550 | 3 | 3 |
| cp-28 | 162-172 | AT550 | 3 | 8 |
| cp-41 | 175-182 | AT550 | 3 | 5 |
| cp-39 | 104-107 | FAM | 4 | 4 |
| cp-51 | 139-144 | FAM | 4 | 6 |
| cp-48 | 154-155 | FAM | 4 | 2 |
| cp-49 | 179-182 | FAM | 4 | 4 |

**Table S5: Characteristics of the mtDNA variants obtained from RADseq data (mtRAD)**

We detected polymorphism on seven mtRADs among ten (numbered 1 to 10 [RAD-loci ID] according to their position on the *N. clarinerva* mitogenome [variant position]). When polymorphism was observed on successive variable nucleotides (di- or tri-nucleotide motifs), we treated these sites as a same locus since possibly due to a single mutation event (i.e. GA-TC and GAA-TTC inversions). Eleven variants were recovered from polymorphic mitochondrial loci. Variant position = Position of the first variable nucleotide on the *N. clarinerva* mitogenome; the polymorphic motif and number of alleles of each locus are also given.

| RAD-loci ID | Variant position | Motif | Number of alleles |
|---|---|---|---|
| mt-2 | 61173 | TTC / GAA | 2 |
| mt-3 | 61262 | A / C | 2 |
| mt-3 | 61715 | C / A | 2 |
| mt-3 | 62034 | T / G | 2 |
| mt-4 | 336941 | GA / TC | 2 |
| mt-5 | 343571 | G / T | 2 |
| mt-5 | 344347 | A / G | 2 |
| mt-6 | 405658 | A / T | 2 |
| mt-7 | 431714 | C / G | 2 |
| mt-8 | 439234 | GA / TC | 2 |
| mt-10 | 576108 | GA / TC - GAA / TTC | 3 |

##### Table S6: Organellar $F_{ST}$ among sampling sites

Estimates of Nei's weighted $F_{ST}$ (Nei, 1973) among forests (upper-right section) and associated *P*-values (lower-left section) inferred from chloroplast microsatellite data (cpSSR), mtDNA sequences (mtRAD + mtSSR) and combined mtDNA and cpSSR data (cp + mt). $F_{ST}$ were not estimated for Ampondrabe where a single *N. spinifolia* individual was found and genotyped. Heat colors, from dark green to dark red (upper-right section), illustrate the differentiation level between pairs of populations (from low to high values, respectively). The significance of *P*-value (for a non-null $F_{ST}$ value) is also highlighted by a color (yellow for $P < 0.05$; brown for $P < 0.01$).

| Forest | Organellar marker(s) | Forest | | | | | | |
|---|---|---|---|---|---|---|---|---|
| | | 1 | 2 | 3 | 4 | 5 | 6 | 7 |
| 1 Ambilondamba | cpSSR | | 0.181 | 0.036 | 0.152 | 0.104 | 0.302 | 0.393 |
| | mtDNA | | 0.302 | 0.173 | 0.327 | 0.254 | 0.532 | 0.284 |
| | cp + mt | | 0.231 | 0.078 | 0.235 | 0.150 | 0.399 | 0.401 |
| 3 Antsiasia | cpSSR | 0.009 | | 0.071 | 0.131 | 0.091 | 0.201 | 0.177 |
| | mtDNA | 0.009 | | 0.199 | 0.450 | 0.269 | 0.613 | 0.292 |
| | cp + mt | 0.012 | | 0.105 | 0.271 | 0.123 | 0.371 | 0.198 |
| 3 Bekaraoka | cpSSR | 0.613 | 0.247 | | 0.106 | 0.040 | 0.210 | 0.152 |
| | mtDNA | 0.033 | 0.030 | | 0.499 | 0.020 | 0.645 | 0.005 |
| | cp + mt | 0.159 | 0.117 | | 0.261 | 0.037 | 0.381 | 0.149 |
| 4 Benanofy | cpSSR | 0.017 | 0.048 | 0.121 | | 0.134 | 0.133 | 0.258 |
| | mtDNA | 0.002 | 0.003 | 0.005 | | 0.489 | 0.058 | 0.415 |
| | cp + mt | 0.015 | 0.009 | 0.014 | | 0.271 | 0.101 | 0.321 |
| 5 Bobankora | cpSSR | 0.102 | 0.189 | 0.789 | 0.083 | | 0.224 | 0.178 |
| | mtDNA | 0.254 | 0.269 | 0.020 | 0.489 | | 0.661 | 0.021 |
| | cp + mt | 0.150 | 0.123 | 0.037 | 0.271 | | 0.392 | 0.164 |
| 6 Solaniampilana | cpSSR | 0.302 | 0.201 | 0.210 | 0.133 | 0.224 | | 0.312 |
| | mtDNA | 0.532 | 0.613 | 0.645 | 0.058 | 0.661 | | 0.615 |
| | cp + mt | 0.399 | 0.371 | 0.381 | 0.101 | 0.392 | | 0.427 |
| 7 Binara | cpSSR | 0.001 | 0.120 | 0.208 | 0.028 | 0.237 | 0.038 | |
| | mtDNA | 0.006 | 0.016 | 0.989 | 0.003 | 0.872 | 0.024 | |
| | cp + mt | 0.003 | 0.021 | 0.079 | 0.005 | 0.119 | 0.015 | |

#### Table S7: Organellar Nei and Edward's *D* among sampling sites

Estimates of Nei's weighted $F_{ST}$ (Nei, 1973) [upper section], Nei's *D* (Nei, 1972) [middle section], and Edward's *D* (Edwards, 1971) [lower section], among forests inferred from mtDNA sequences (mtRAD + mtSSR, left panel), chloroplast microsatellite data (cpSSR, middle panel), and combined mtDNA and cpSSR data (cp + mt, right panel). Distances and differentiation metrics were not estimated for Ampondrabe where a single *N. spinifolia* individual was found and genotyped. Heat colors, from dark green to dark red, illustrate the differentiation level between pairs of populations (from low to high values, respectively).

**Nei's Fst**

| mtDNA | Ambilondamba | Antsiasia | Bekaraoka | Benanofy | Binara | Bobankora | Solaniampilana |
|---|---|---|---|---|---|---|---|
| Ambilondamba | 0.000 | 0.302 | 0.173 | 0.327 | 0.284 | 0.254 | 0.532 |
| Antsiasia | 0.302 | 0.000 | 0.199 | 0.450 | 0.292 | 0.269 | 0.613 |
| Bekaraoka | 0.173 | 0.199 | 0.000 | 0.499 | 0.005 | 0.020 | 0.645 |
| Benanofy | 0.327 | 0.450 | 0.499 | 0.000 | 0.415 | 0.489 | 0.058 |
| Binara | 0.284 | 0.292 | 0.005 | 0.415 | 0.000 | 0.021 | 0.615 |
| Bobankora | 0.254 | 0.269 | 0.020 | 0.489 | 0.021 | 0.000 | 0.661 |
| Solaniampilana | 0.532 | 0.613 | 0.645 | 0.058 | 0.615 | 0.661 | 0.000 |

| cpSSR | Ambilondamba | Antsiasia | Bekaraoka | Benanofy | Binara | Bobankora | Solaniampilana |
|---|---|---|---|---|---|---|---|
| Ambilondamba | 0.000 | 0.193 | 0.041 | 0.152 | 0.421 | 0.117 | 0.301 |
| Antsiasia | 0.193 | 0.000 | 0.071 | 0.131 | 0.177 | 0.091 | 0.201 |
| Bekaraoka | 0.041 | 0.071 | 0.000 | 0.106 | 0.150 | 0.039 | 0.208 |
| Benanofy | 0.152 | 0.131 | 0.106 | 0.000 | 0.258 | 0.134 | 0.133 |
| Binara | 0.421 | 0.177 | 0.150 | 0.258 | 0.000 | 0.178 | 0.312 |
| Bobankora | 0.117 | 0.091 | 0.039 | 0.134 | 0.178 | 0.000 | 0.224 |
| Solaniampilana | 0.301 | 0.201 | 0.208 | 0.133 | 0.312 | 0.224 | 0.000 |

| cp + mt | Ambilondamba | Antsiasia | Bekaraoka | Benanofy | Binara | Bobankora | Solaniampilana |
|---|---|---|---|---|---|---|---|
| Ambilondamba | 0.000 | 0.231 | 0.078 | 0.235 | 0.401 | 0.150 | 0.399 |
| Antsiasia | 0.231 | 0.000 | 0.105 | 0.271 | 0.198 | 0.123 | 0.371 |
| Bekaraoka | 0.078 | 0.105 | 0.000 | 0.261 | 0.149 | 0.037 | 0.381 |
| Benanofy | 0.235 | 0.271 | 0.261 | 0.000 | 0.321 | 0.271 | 0.101 |
| Binara | 0.401 | 0.198 | 0.149 | 0.321 | 0.000 | 0.164 | 0.427 |
| Bobankora | 0.150 | 0.123 | 0.037 | 0.271 | 0.164 | 0.000 | 0.392 |
| Solaniampilana | 0.399 | 0.371 | 0.381 | 0.101 | 0.427 | 0.392 | 0.000 |

**Nei's D**

| mtDNA | Ambilondamba | Antsiasia | Bekaraoka | Benanofy | Binara | Bobankora | Solaniampilana |
|---|---|---|---|---|---|---|---|
| Ambilondamba | 0.000 | 0.115 | 0.062 | 0.307 | 0.064 | 0.067 | 0.530 |
| Antsiasia | 0.115 | 0.000 | 0.068 | 0.529 | 0.071 | 0.070 | 0.771 |
| Bekaraoka | 0.062 | 0.068 | 0.000 | 0.397 | 0.001 | 0.003 | 0.620 |
| Benanofy | 0.307 | 0.529 | 0.397 | 0.000 | 0.392 | 0.392 | 0.029 |
| Binara | 0.064 | 0.071 | 0.001 | 0.392 | 0.000 | 0.003 | 0.606 |
| Bobankora | 0.067 | 0.070 | 0.003 | 0.392 | 0.003 | 0.000 | 0.604 |
| Solaniampilana | 0.530 | 0.771 | 0.620 | 0.029 | 0.606 | 0.604 | 0.000 |

| cpSSR | Ambilondamba | Antsiasia | Bekaraoka | Benanofy | Binara | Bobankora | Solaniampilana |
|---|---|---|---|---|---|---|---|
| Ambilondamba | 0.000 | 0.223 | 0.060 | 0.168 | 0.469 | 0.123 | 0.387 |
| Antsiasia | 0.223 | 0.000 | 0.158 | 0.235 | 0.267 | 0.155 | 0.371 |
| Bekaraoka | 0.060 | 0.158 | 0.000 | 0.161 | 0.361 | 0.054 | 0.389 |
| Benanofy | 0.168 | 0.235 | 0.161 | 0.000 | 0.463 | 0.188 | 0.164 |
| Binara | 0.469 | 0.267 | 0.361 | 0.463 | 0.000 | 0.273 | 0.535 |
| Bobankora | 0.123 | 0.155 | 0.054 | 0.188 | 0.273 | 0.000 | 0.347 |
| Solaniampilana | 0.387 | 0.371 | 0.389 | 0.164 | 0.535 | 0.347 | 0.000 |

| cp + mt | Ambilondamba | Antsiasia | Bekaraoka | Benanofy | Binara | Bobankora | Solaniampilana |
|---|---|---|---|---|---|---|---|
| Ambilondamba | 0.000 | 0.175 | 0.067 | 0.238 | 0.255 | 0.100 | 0.475 |
| Antsiasia | 0.175 | 0.000 | 0.109 | 0.400 | 0.156 | 0.106 | 0.595 |
| Bekaraoka | 0.067 | 0.109 | 0.000 | 0.283 | 0.155 | 0.024 | 0.524 |
| Benanofy | 0.238 | 0.400 | 0.283 | 0.000 | 0.431 | 0.296 | 0.086 |
| Binara | 0.255 | 0.156 | 0.155 | 0.431 | 0.000 | 0.122 | 0.608 |
| Bobankora | 0.100 | 0.106 | 0.024 | 0.296 | 0.122 | 0.000 | 0.505 |
| Solaniampilana | 0.475 | 0.595 | 0.524 | 0.086 | 0.608 | 0.505 | 0.000 |

**Edward's D**

| mtDNA | Ambilondamba | Antsiasia | Bekaraoka | Benanofy | Binara | Bobankora | Solaniampilana |
|---|---|---|---|---|---|---|---|
| Ambilondamba | 0.000 | 0.302 | 0.245 | 0.447 | 0.246 | 0.260 | 0.563 |
| Antsiasia | 0.302 | 0.000 | 0.231 | 0.568 | 0.258 | 0.244 | 0.655 |
| Bekaraoka | 0.245 | 0.231 | 0.000 | 0.503 | 0.093 | 0.097 | 0.603 |
| Benanofy | 0.447 | 0.568 | 0.503 | 0.000 | 0.508 | 0.513 | 0.184 |
| Binara | 0.246 | 0.258 | 0.093 | 0.508 | 0.000 | 0.086 | 0.607 |
| Bobankora | 0.260 | 0.244 | 0.097 | 0.513 | 0.086 | 0.000 | 0.609 |
| Solaniampilana | 0.563 | 0.655 | 0.603 | 0.184 | 0.607 | 0.609 | 0.000 |

| cpSSR | Ambilondamba | Antsiasia | Bekaraoka | Benanofy | Binara | Bobankora | Solaniampilana |
|---|---|---|---|---|---|---|---|
| Ambilondamba | 0.000 | 0.482 | 0.315 | 0.415 | 0.638 | 0.390 | 0.551 |
| Antsiasia | 0.482 | 0.000 | 0.402 | 0.493 | 0.543 | 0.427 | 0.582 |
| Bekaraoka | 0.315 | 0.402 | 0.000 | 0.400 | 0.548 | 0.265 | 0.544 |
| Benanofy | 0.415 | 0.493 | 0.400 | 0.000 | 0.614 | 0.437 | 0.423 |
| Binara | 0.638 | 0.543 | 0.548 | 0.614 | 0.000 | 0.509 | 0.628 |
| Bobankora | 0.390 | 0.427 | 0.265 | 0.437 | 0.509 | 0.000 | 0.554 |
| Solaniampilana | 0.551 | 0.582 | 0.544 | 0.423 | 0.628 | 0.554 | 0.000 |

| cp + mt | Ambilondamba | Antsiasia | Bekaraoka | Benanofy | Binara | Bobankora | Solaniampilana |
|---|---|---|---|---|---|---|---|
| Ambilondamba | 0.000 | 0.415 | 0.286 | 0.437 | 0.501 | 0.341 | 0.563 |
| Antsiasia | 0.415 | 0.000 | 0.342 | 0.538 | 0.435 | 0.358 | 0.623 |
| Bekaraoka | 0.286 | 0.342 | 0.000 | 0.454 | 0.405 | 0.200 | 0.579 |
| Benanofy | 0.437 | 0.538 | 0.454 | 0.000 | 0.569 | 0.479 | 0.326 |
| Binara | 0.501 | 0.435 | 0.405 | 0.569 | 0.000 | 0.381 | 0.629 |
| Bobankora | 0.341 | 0.358 | 0.200 | 0.479 | 0.381 | 0.000 | 0.590 |
| Solaniampilana | 0.563 | 0.623 | 0.579 | 0.326 | 0.629 | 0.590 | 0.000 |

**Table S8: Nuclear genetic differentiation among *Noronhia spinifolia* sampling sites**

Estimates of Reynolds' weighted $F_{ST}$ (Reynolds et al., 1983) (upper-right section) and associated $P$-values (lower-left section), among forests, inferred from genotype likelihood in ANGSD. $F_{ST}$ were not estimated for Ampondrabe where a single *N. spinifolia* individual was found and genotyped. Heat colors, from dark green to dark red, illustrate the differentiation level between pairs of populations (from low to high values, respectively). The significance of $P$-value (for a non-null $F_{ST}$ value) is also highlighted by a color (yellow for $P < 0.05$; brown for $P < 0.01$).

| Population | 1 | 2 | 3 | 4 | 5 | 6 | 7 |
|---|---|---|---|---|---|---|---|
| 1 Ambilondamba | | 0.112 | 0.170 | 0.149 | 0.119 | 0.136 | 0.138 |
| 2 Antsiasia | 0.297 | | 0.133 | 0.147 | 0.128 | 0.112 | 0.134 |
| 3 Bekaraoka | <0.01 | <0.01 | | 0.132 | 0.210 | 0.089 | 0.108 |
| 4 Benanofy | <0.01 | 0.010 | <0.01 | | 0.179 | 0.124 | 0.115 |
| 5 Binara | 0.119 | 0.020 | <0.01 | <0.01 | | 0.139 | 0.172 |
| 6 Bobankora | 0.020 | 0.138 | <0.01 | 0.011 | <0.01 | | 0.111 |
| 7 Solaniampilana | <0.01 | <0.01 | <0.01 | 0.010 | <0.01 | 0.011 | |

##### Table S9: Organellar microsatellite genotypes

*Also available in bioRxiv:* https://www.biorxiv.org/content/10.1101/2020.11.25.394544v4
Sample ID: Sample identifier; Forest: local name of the forest; Latitude and Longitude are given in WGS 1984, degree decimal; Marker ID: organellar microsatellite marker identifier, all markers are located on the plastome except mt-32, which is located on the mitochondrial genome. Fifteen chloroplast microsatellites loci (cpSSRs) plus one mitochondrial SSR locus (mt-32) were amplified in four multiplex reactions (Table S2), each with a distinct fluorochrome (YAK, AT565, AT550, FAM). For each marker and individual, the allele size is reported in the table.

##### Table S10: Mitochondrial RADseq genotypes

*Also available in bioRxiv:* https://www.biorxiv.org/content/10.1101/2020.11.25.394544v4
Sample ID: Sample identifier; Forest: local name of the forest; Latitude and Longitude are given in WGS 1984, degree decimal; Marker ID: alignment coordinates on *N. clarinerva* mitogenome. Original variants: SNPs called from alignment files; Summarized variants: summary of original variants considering simplifying successive variable nucleotides (di- or tri-nucleotide motifs) for analytical purposes.

#### Supporting figures

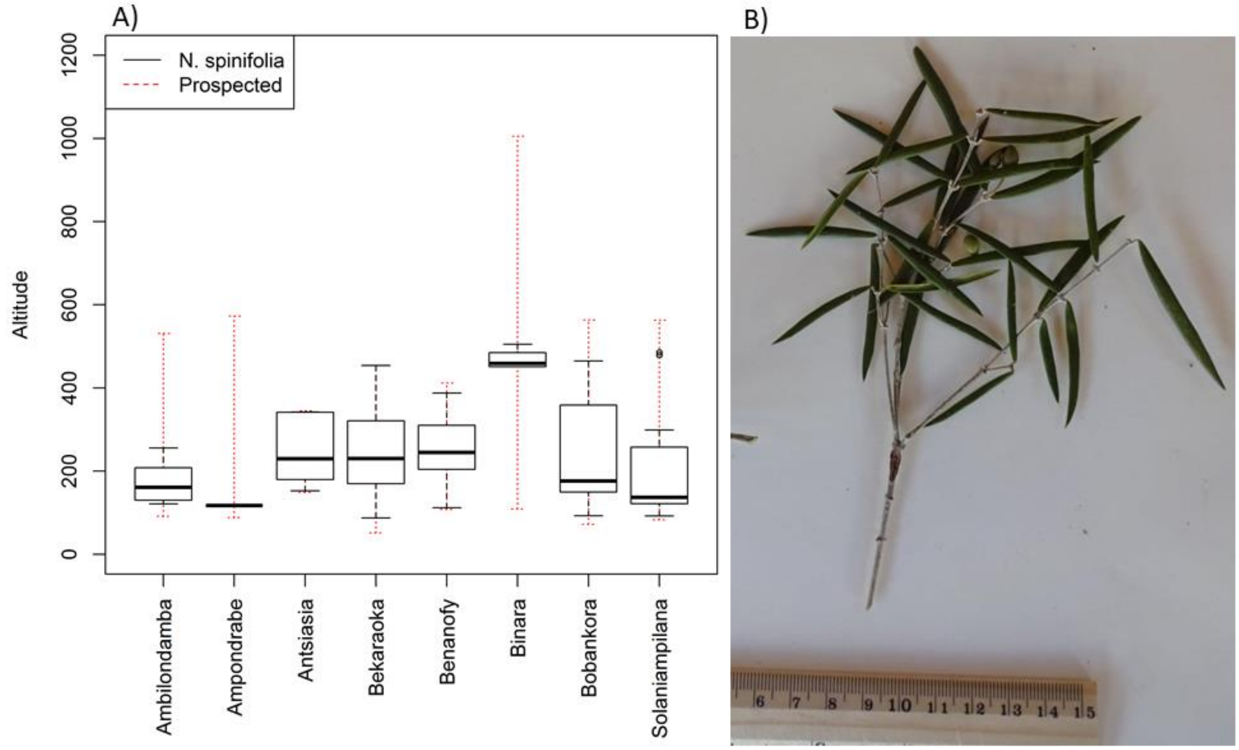


**Figure S1: Altitudinal distribution range of *Noronhia spinifolia* in the Loky-Manambato region**

In **A)**, the boxplot of sampled trees by forest fragment illustrates the altitudinal range at which the species occurs in the Loky-Manambato (LM) region. The red dotted range illustrates the altitudinal range prospected in each forest. It is worth noting that the species occurs at a higher elevation (451-505 m) in Binara. This unexpected pattern, repeatedly found in other Malagasy olives species of the LM region (*N. ankaranensis*, *N. candicans*, *N. christenseniana*, and *N. oblanceolata*; JS & GB unpublished data), suggests that particular processes (e.g. Pleistocene and Holocene climatic shifts) at stake in the region need to be investigated with greater care. The picture in **B**) illustrates a branch of *N. spinifolia* with its easily recognizable spiny leaves and carrying three unripe fruits.

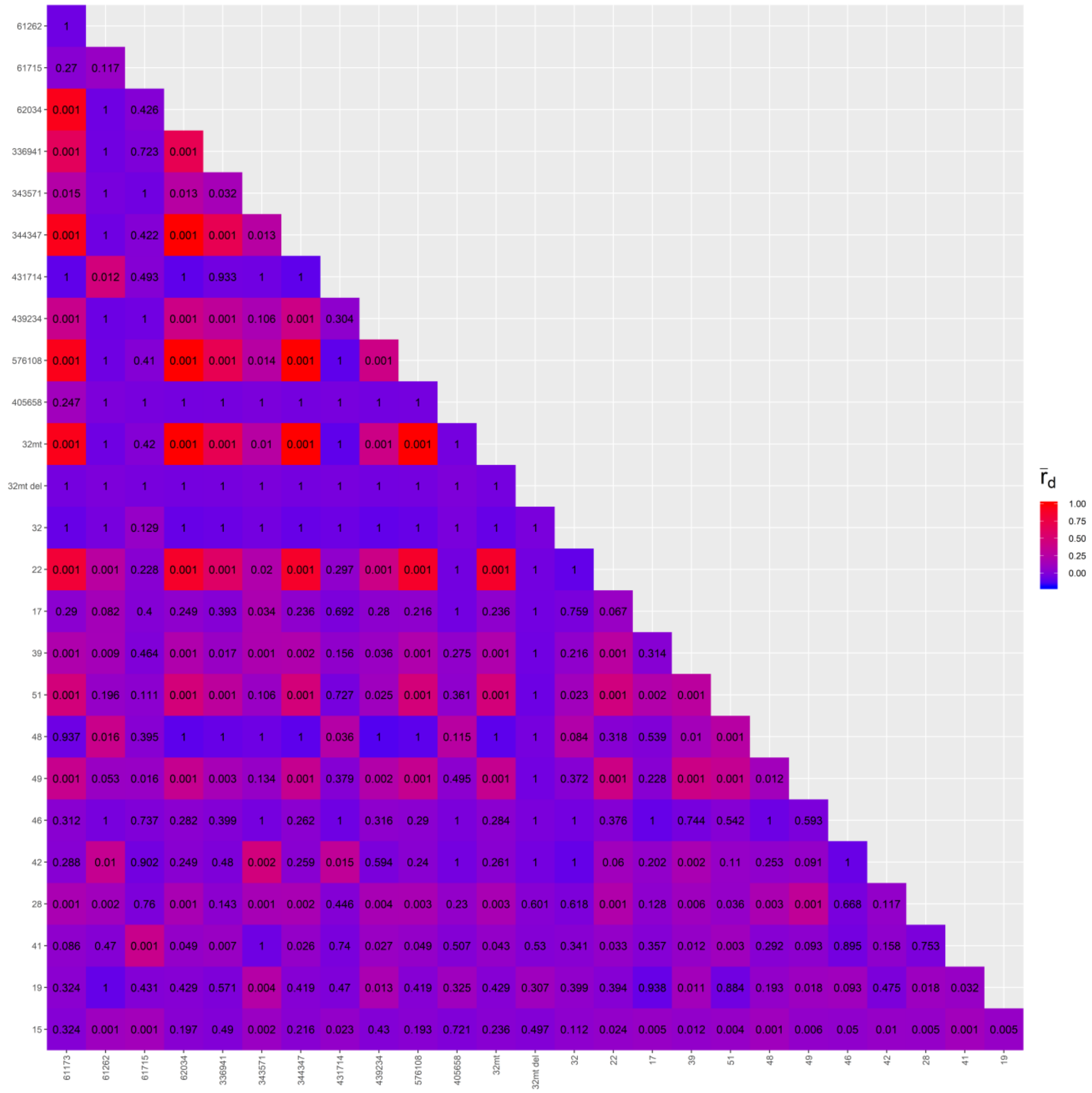


#### Figure S2: Linkage disequilibrium in organellar DNA data

The Agapow & Burt $r_d$ values of linkage disequilibrium among organellar (mitochondrial and chloroplast) loci are represented with the red to blue color scale, their associated *p*-values are noted in their respective cells, x and y axis labels represent the cpSSR and mtDNA locus ID. A high (and significant) $r_d$ value (>0.4; cf. color scale) indicates a strong linkage between the makers, suggesting they are not recombining. However, low and non-significant LD values are more complex to interpret because these can result from homoplasy, recombination, and/or low diversity at the investigated loci (Mueller, 2004).

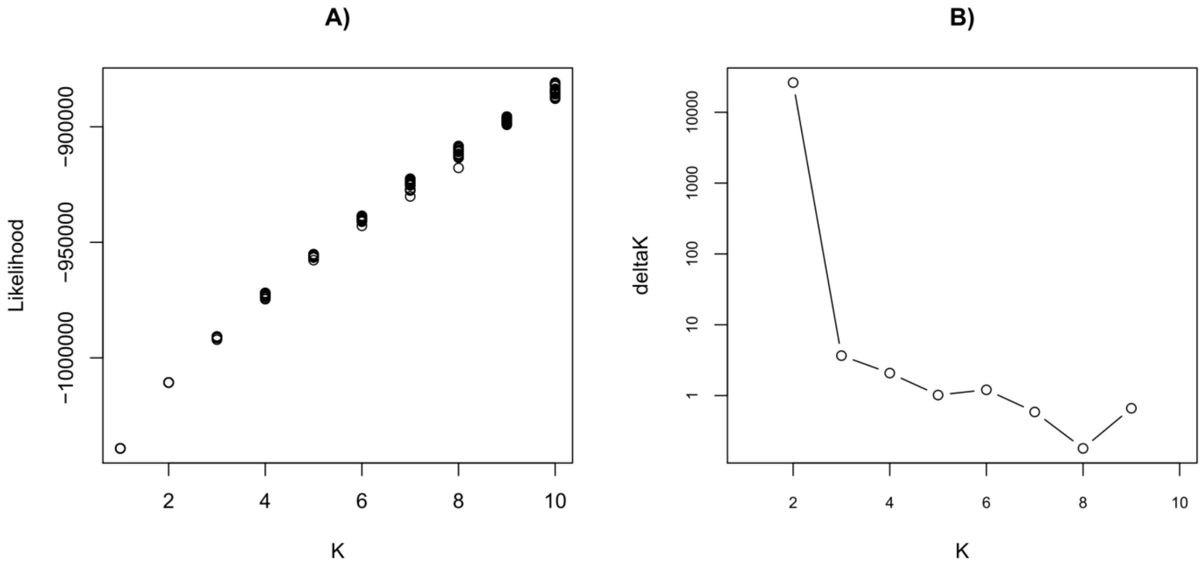


**Figure S3: Number of nuclear genetic clusters best explaining the data when using NgsAdmix**

Likelihood **(A)** and delta*K* **(B)** results. Although the likelihood continuously increases, we find little evidence that *K* values > 1 better explain the data than the results for *K* = 1. Considering the continuous increase and the rather continuous (as opposed to discrete) observed pattern of structure, we chose to present results for *K* = 2, 3 and 4 that provide decent illustrations of how ngsAdmix explains the data.

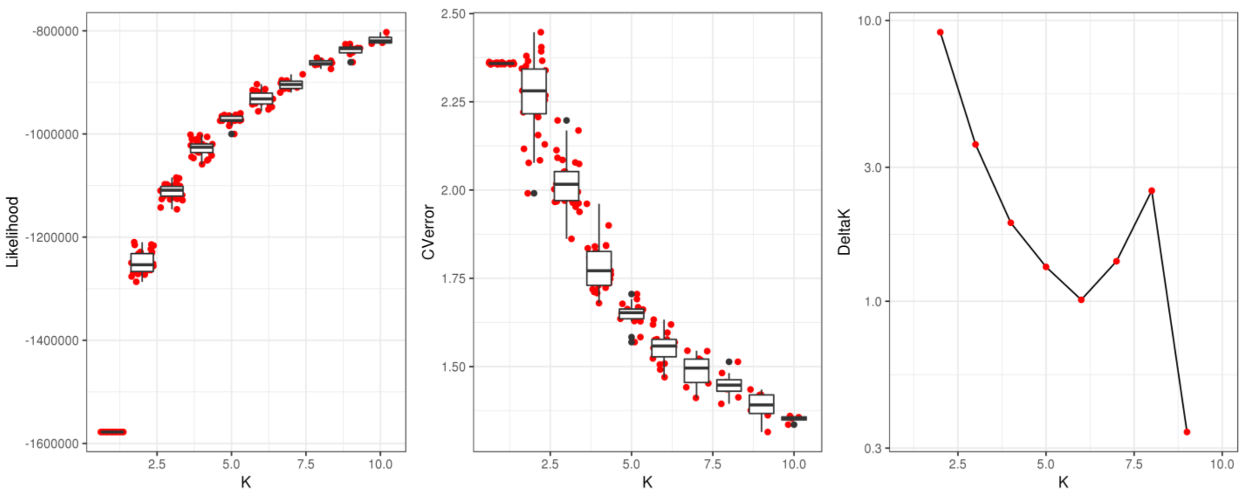


**Figure S4: Number of nuclear genetic clusters best explaining the data when using Admixture**

Likelihood **(left panel)**, cross-validation error **(middle panel)** and delta*K* **(right panel)** results obtained from 20 Admixture runs per *K* value. Although the likelihood (**left panel**) continuously increases, we find little evidence that *K* values > 1 better explain the data than the results for *K* = 1. The cross-validation error (**middle panel**) shows no typical signature of a clear discrete structure. The delta*K* (**right panel**) shows small peaks for *K* = 2 and 8.

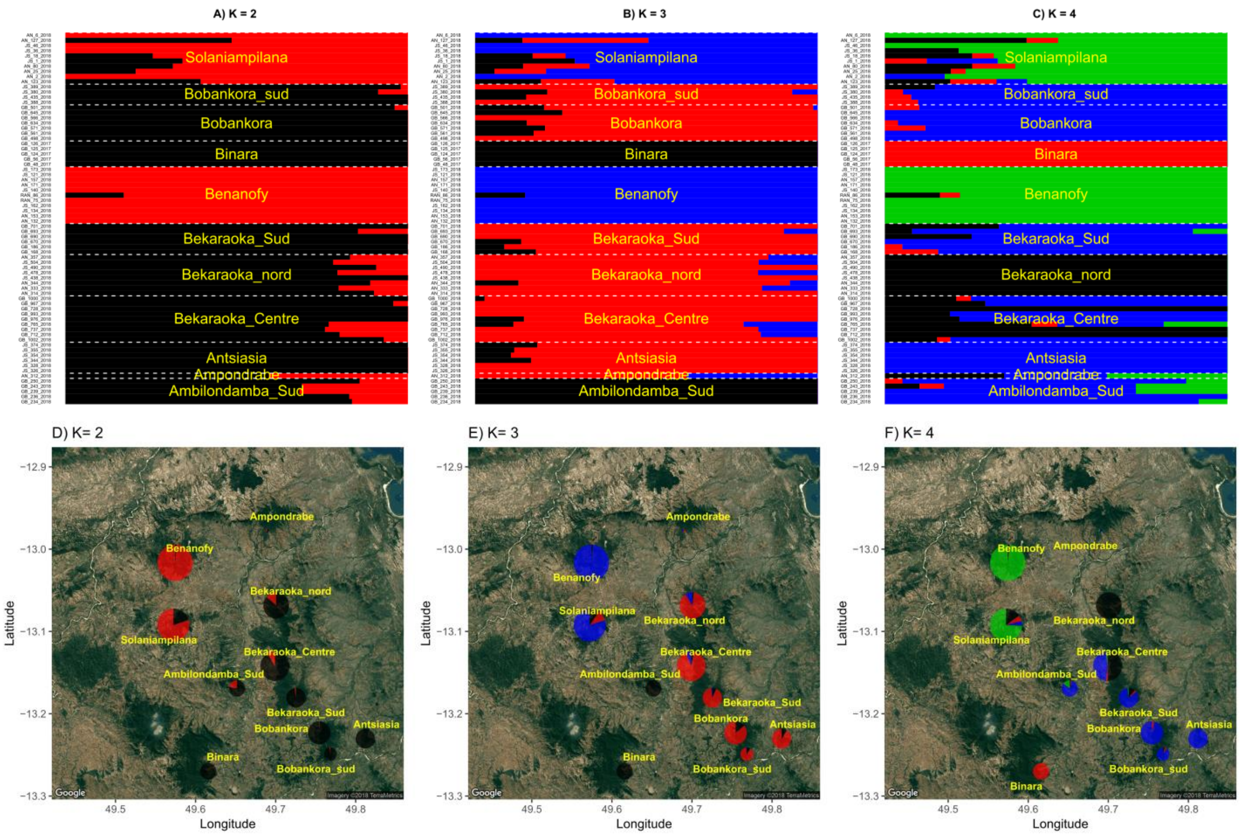


#### Figure S5: Genetic structure in *Noronhia spinifolia*

Individual ancestry proportions inferred in NgsAdmix **(A, B, C)** and geographic representation of ancestry proportions per site **(D, E, F)**. Results are represented for $K$ = 2 **(A, D)**, $K$ = 3 **(B, E)** and $K$ = 4 **(C, F)** according to likelihood and delta$K$ results in Fig. S8. Pie chart size is proportional to the number of samples analyzed per location. Pie shares **(D, E, F)** are proportional to the sums of individual ancestry proportions in **A, B** and **C**. Yellow labels correspond to sampling sites.

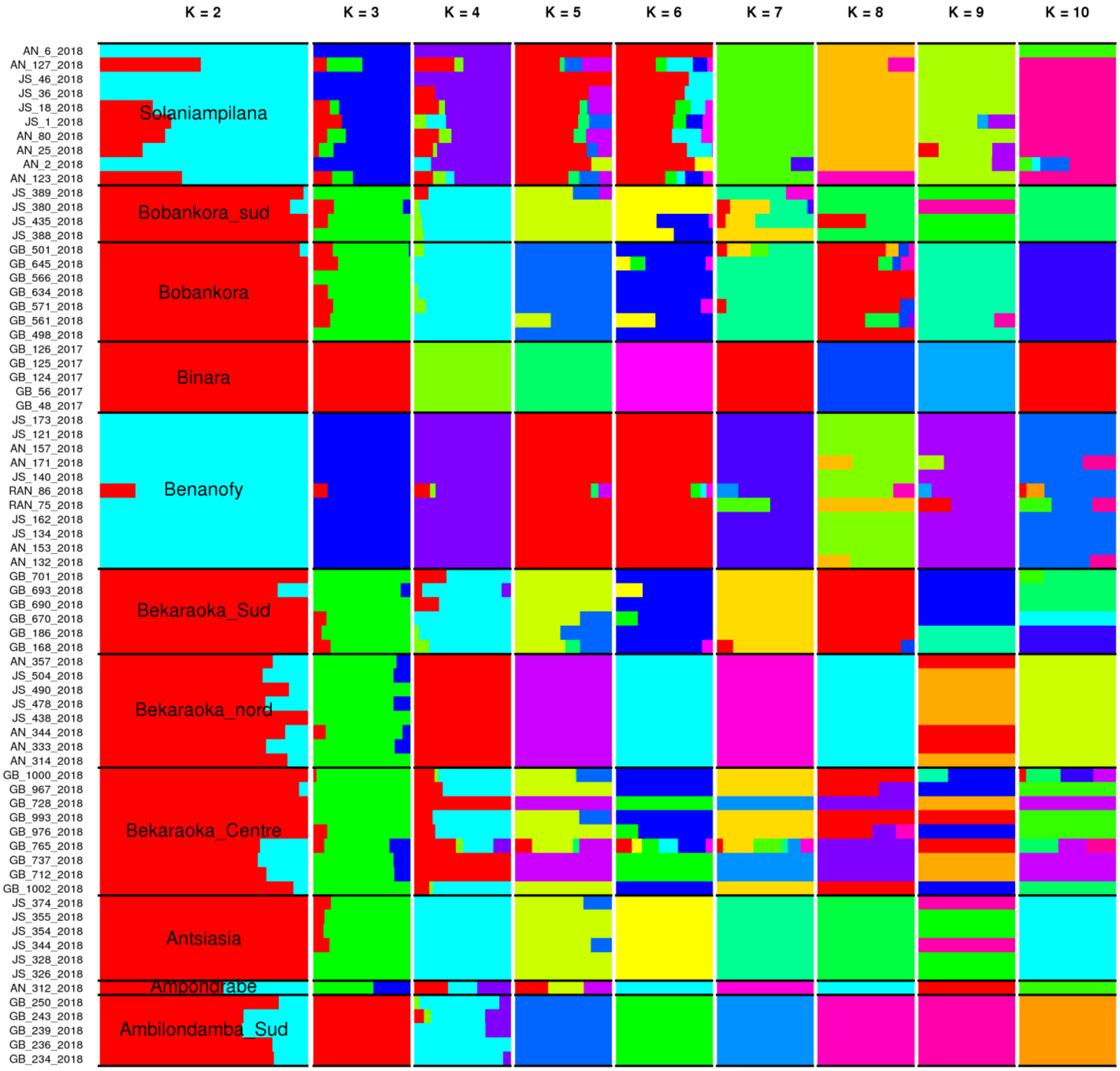


**Figure S6: ngsAdmix ancestry proportion estimates for *K* = 2 to 10**

Individual ancestry proportions inferred in NgsAdmix, for all tested *K* values (2-10). Black labels in *K* = 2 correspond to sampling sites. The figure illustrates that, although likelihood and delta*K* estimates (Fig. S8) did point toward a strong structure in discrete clusters, the NgsAdmix inferences are able to extract geographically coherent signal for all *K* values, which tends to be of a continuous nature (Fig. S10) for low *K* values but tends towards distinguishing almost each sampling site as a discrete cluster for higher *K* values.

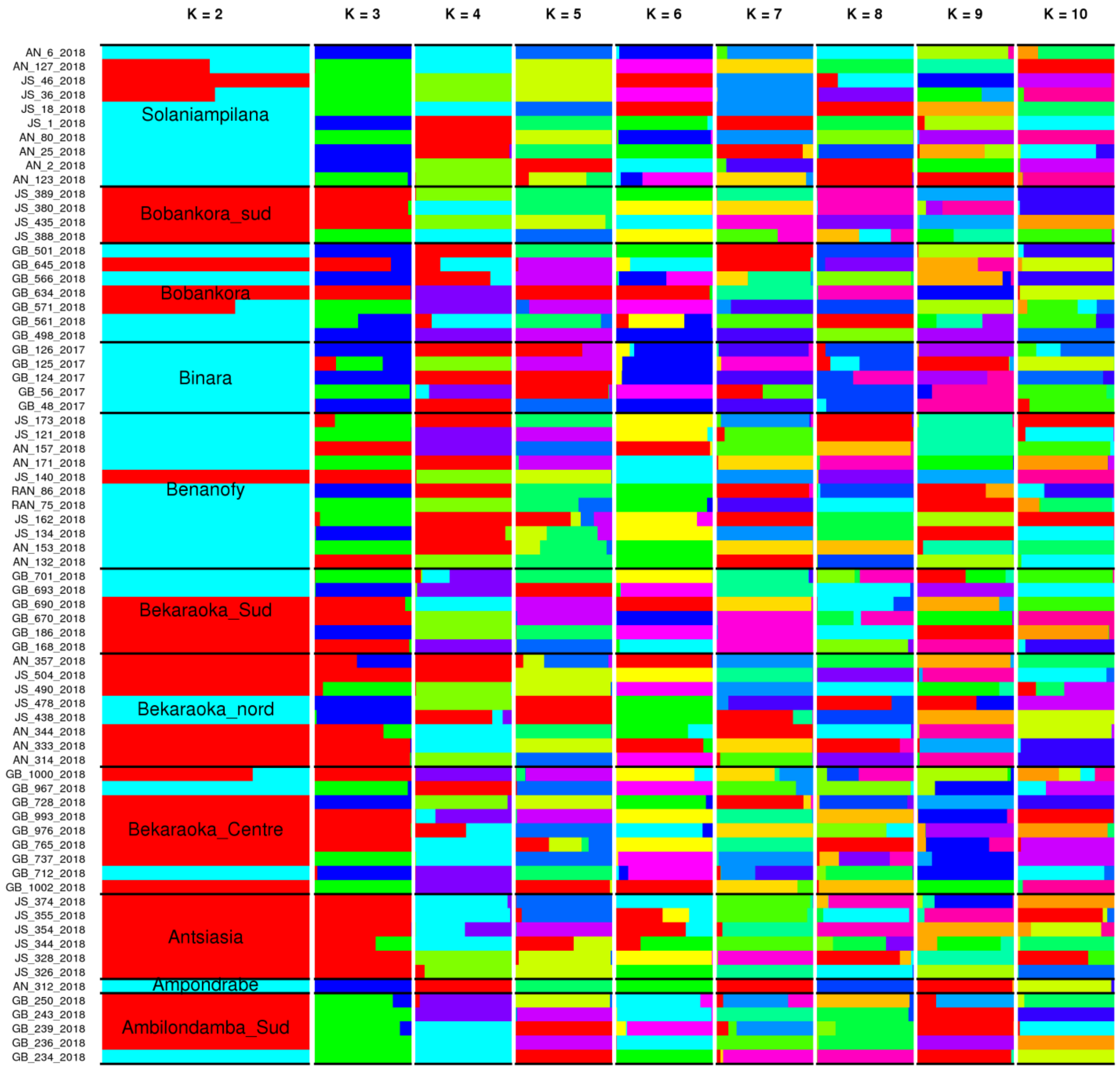


#### Figure S7: Admixture ancestry proportion estimates for *K* = 2 to 10

Individual ancestry proportions inferred in Admixture for all tested *K* values (2-10). Black labels in *K* = 2 correspond to sampling sites. The figure illustrates that, in contrast to NgsAdmix, Admixture could not recover any coherent signal of structure from the nuclear SNPs called with Stacks.

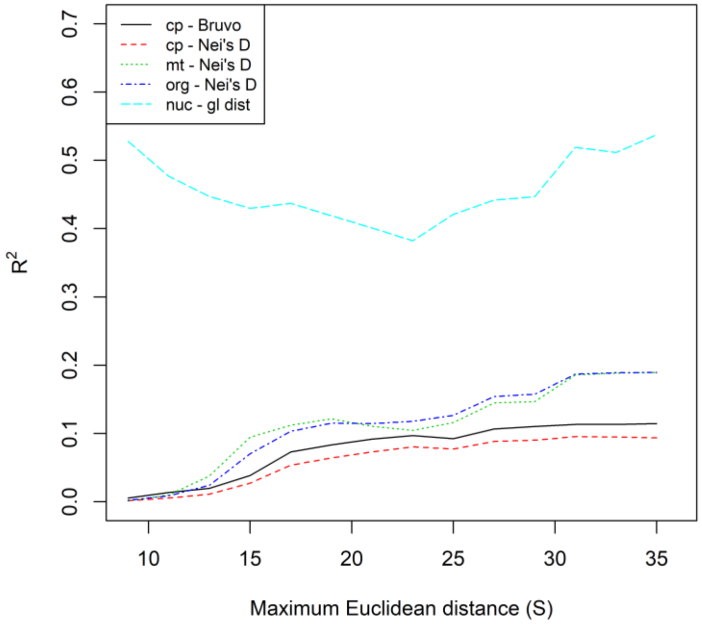


**Figure S8: Geographic scale influence on isolation by distance (IBD)**

Plot of the cumulative variance explained by IBD ($R^2$) against the maximum pairwise geographic Euclidean distance (S) estimated from chloroplast microsatellite data (cp – Bruvo and cp - Nei's D), mitochondrial RAD data (mt – Nei's D), combined organellar DNA polymorphism data (org – Nei's D), and nuclear RAD data (nuc - gl dist). Mantel tests conducted on subsets of pairwise data defined by their maximum geographic distance (S) between samples by 2,000-m increments. For most genetic data and distances, the best fit to the IBD model is obtained using all the data, i.e. at the largest geographic scale. The highest fit is found with the GL covariance matrix estimated in PCAngsd (nuc - gl dist) at a maximum geographic distance (S) of 35 km from nuclear data ($R^2$ = 0.52).

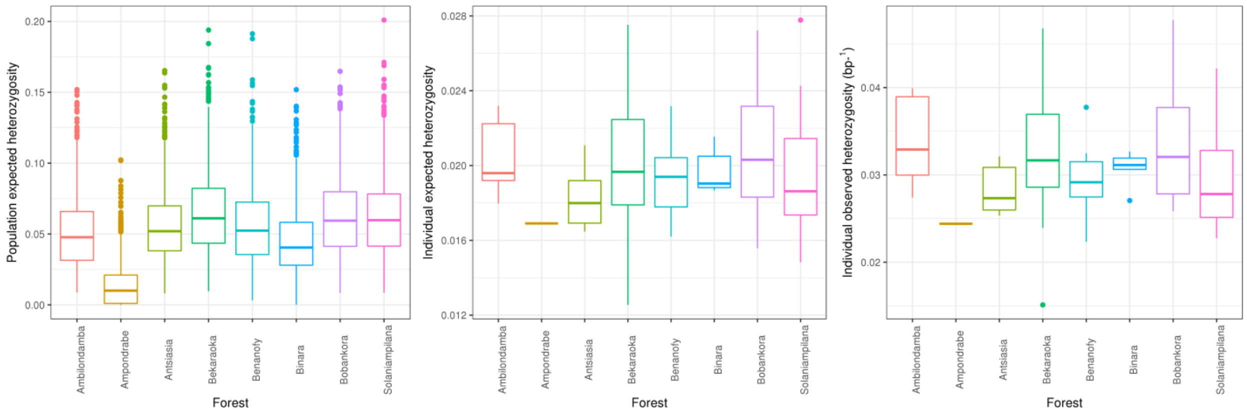


**Figure S9: *Noronhia spinifolia*'s genetic diversity**

Genetic diversity ($H_E$ & $H_O$) across forests. The forest-based (**left panel**) and individual-based (**central panel**) expected heterozygosities were estimated following Fumagalli (2013), whereas the individual-based observed heterozygosities (**right panel**) were estimated as the proportion of heterozygous sites from genotype likelihood (GL) based on folded site frequency estimated in ANGSD (Pedersen et al., 2018). The represented expected heterozygosities were estimated across 1 000bp windows. All three diversity indices are multiplied by ten for representation purposes. Note that Ampondrabe low $H_E$ estimates (left panel) are based on one sample only and may therefore be underestimated.

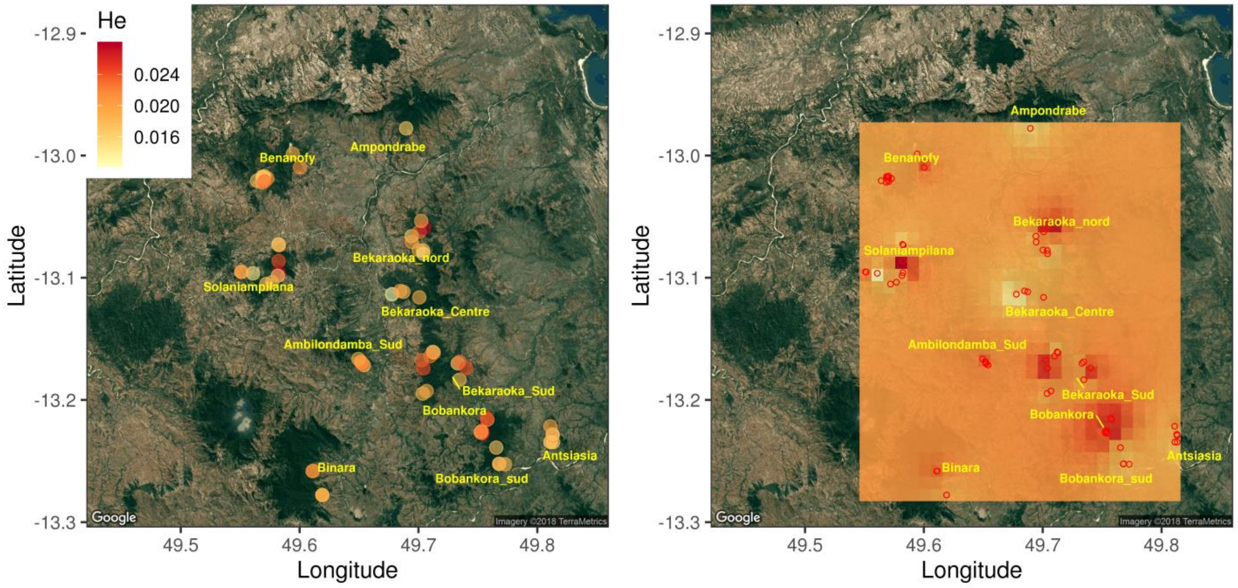


**Figure S10: Spatial distribution of nuclear genetic diversity in *Noronhia spinifolia***

Individual-based expected heterozygosity estimates ($H_E$) are represented on the **left panel** and their inverse distance weighted interpolation is represented on the **right panel**. The figures show that the genetic diversity, estimated following Fumagalli (2013), is not homogeneously distributed in space with areas harboring higher levels of genetic diversity. The geographic inverse distance weighted interpolation of $H_O$ was inferred with the R package *gstat* (Pebesma & Heuvelink, 2016). Expected heterozygosities are multiplied by ten for representation purposes.

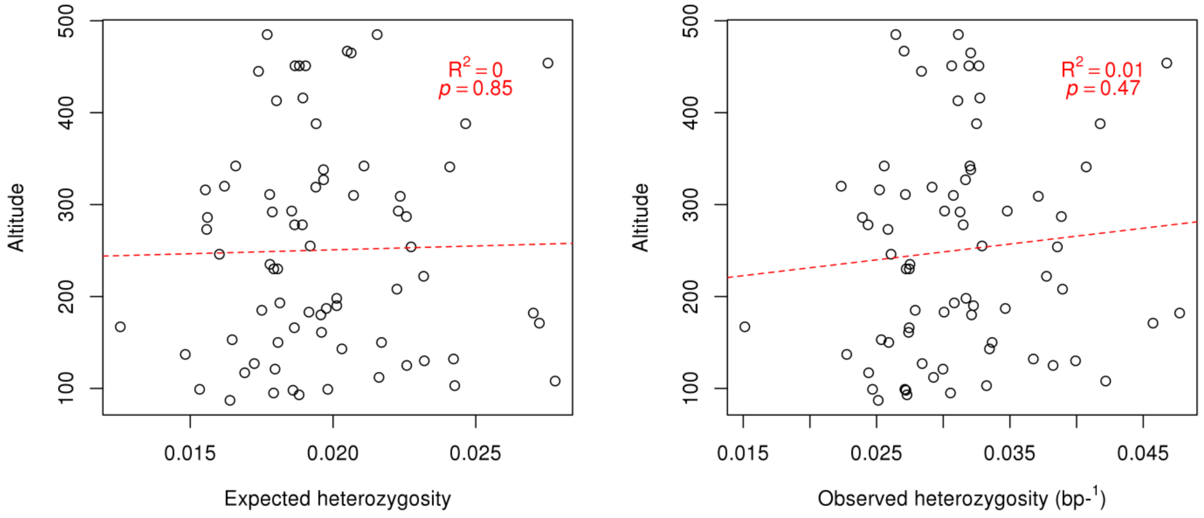


**Figure S11: Altitude effect on *Noronhia spinifolia*'s genetic diversity**

Representation of *N. spinifolia* genetic diversity as a function of elevation. This figure suggests no effect of elevation on nuclear genetic diversity. Individual-based expected heterozygosity ($H_E$), left plot, estimated following Fumagalli (2013) and observed heterozygosity ($H_O$), right plot, estimated following Pedersen et al. (2018). Both diversity indices are multiplied by ten for representation purposes.

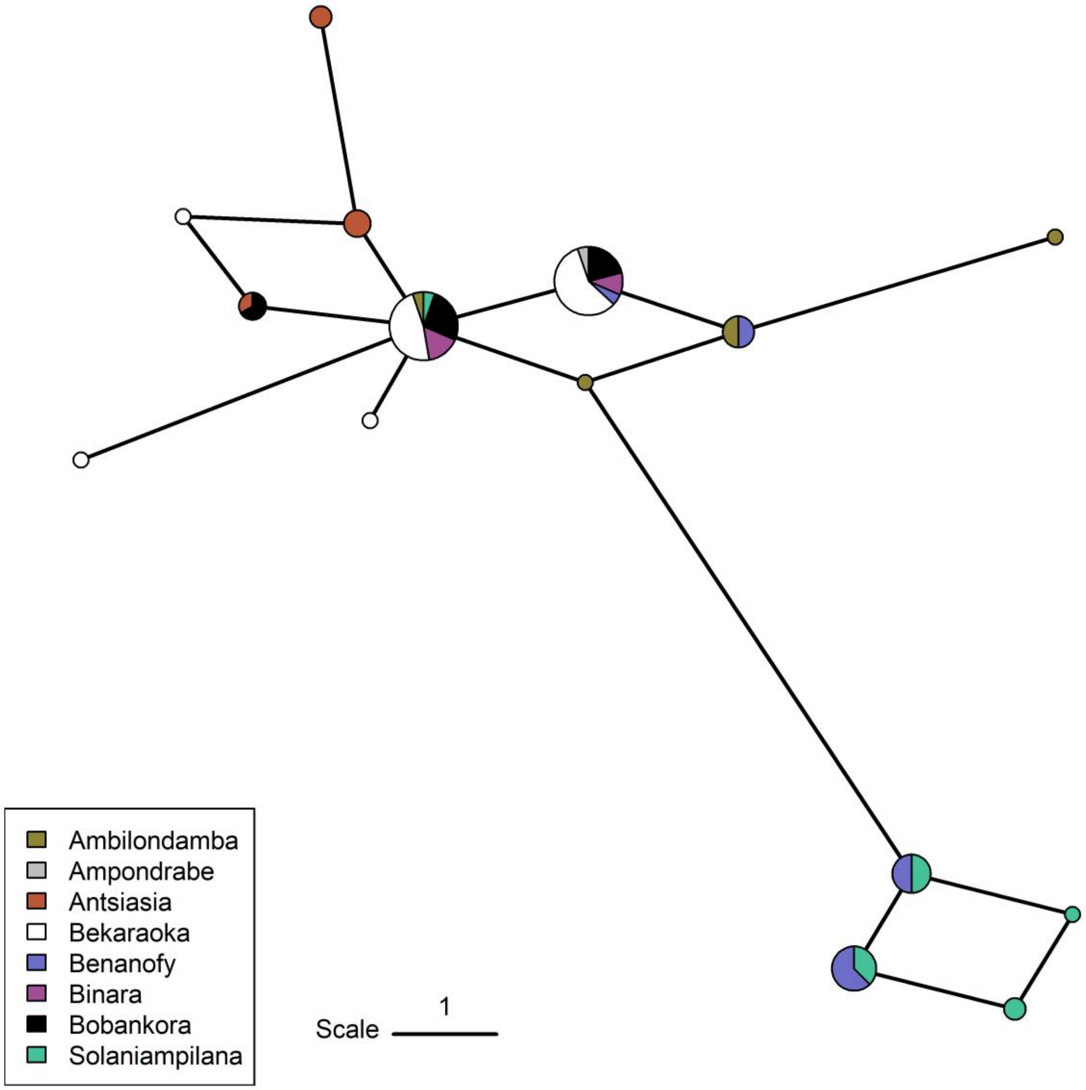


**Figure S12: *Noronhia spinifolia* mtDNA haplotype network**

Branch length is proportional to the Manhattan distance between mitotypes (divided by 2; i.e. equivalent to the number of mutations between haplotypes). Pie chart size is proportional to the number of accessions for a given mitotype. All edges of equal weight are represented. The network shows some spatial structure, with, for instance, some haplotypes from Solaniampilana and Benanofy grouping together at the bottom of the network.

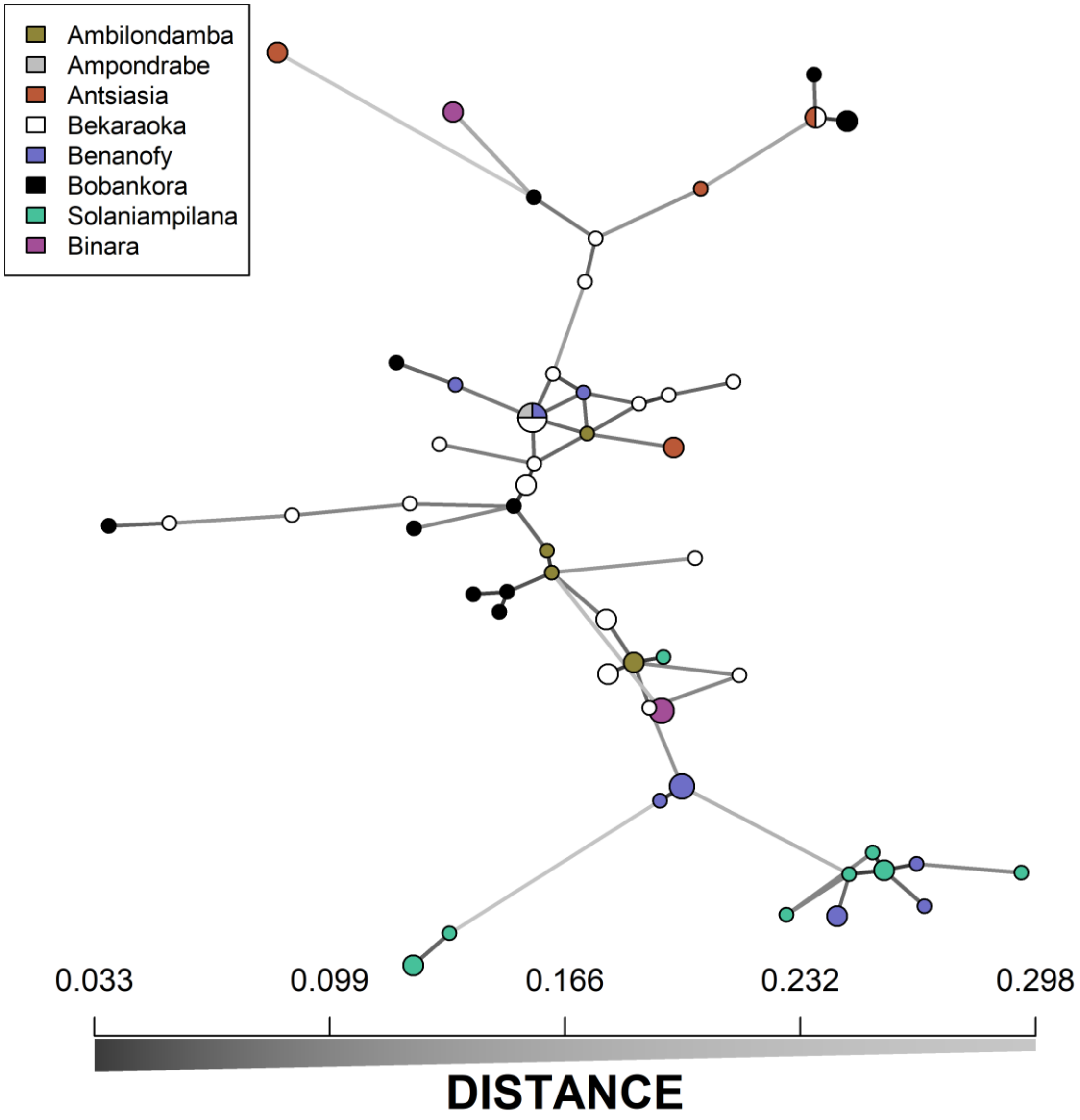


**Figure S13: *Noronhia spinifolia* chlorotype network**

Branch length and gray scale are proportional to the Bruvo's genetic distance between chlorotypes. Pie chart size is proportional to the number of accessions with a given chlorotype. All edges of equal weight are represented. The network highlights large chloroplast diversity with only two haplotypes shared by individuals from at least two forests. It further shows a limited spatial structure, with, for instance, some haplotypes from Solaniampilana and Benanofy grouping together at the bottom of the network.

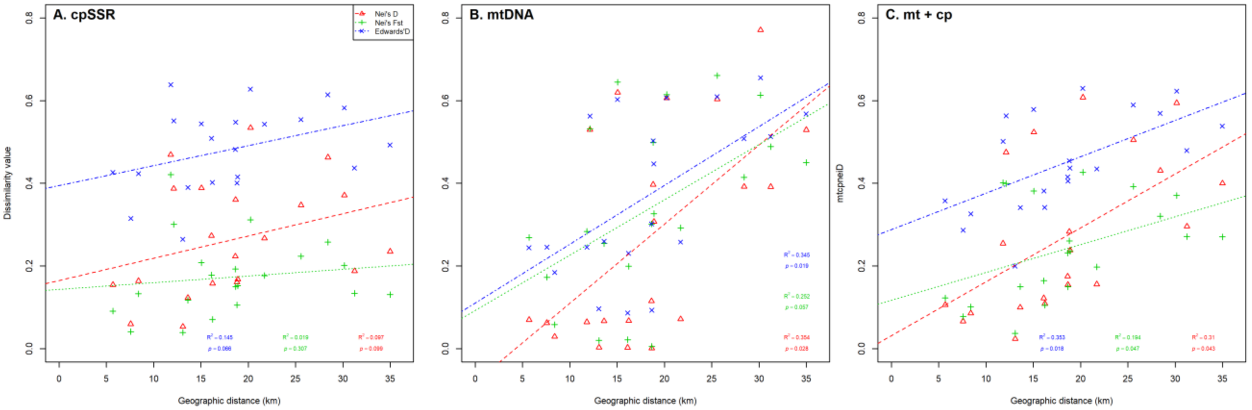


**Figure S14: Forest based organellar isolation by distance in *Noronhia spinifolia***

Graphic representation of the relationship between geographic and genetic distances (isolation by distance). The forest-based genetic distances estimated from chloroplast microsatellite data **(A),** mitochondrial data **(B)**, and combined organellar DNA data **(C)**. The linear model (dashed lines), the Mantel tests explained variance ($R^2$) and $p$-value ($p$) are shown to support the represented trends. The figure shows a non-significant positive relationship for chloroplast data, significant for mitochondrial and combined organellar data with Euclidean geographic distances suggesting that the organellar gene flow (seed dispersal) may be impacted by geographic features. The figure further shows an overall stronger relationship for Nei's $D$ and Edward's $D$ than for the $F_{ST}$ metric.

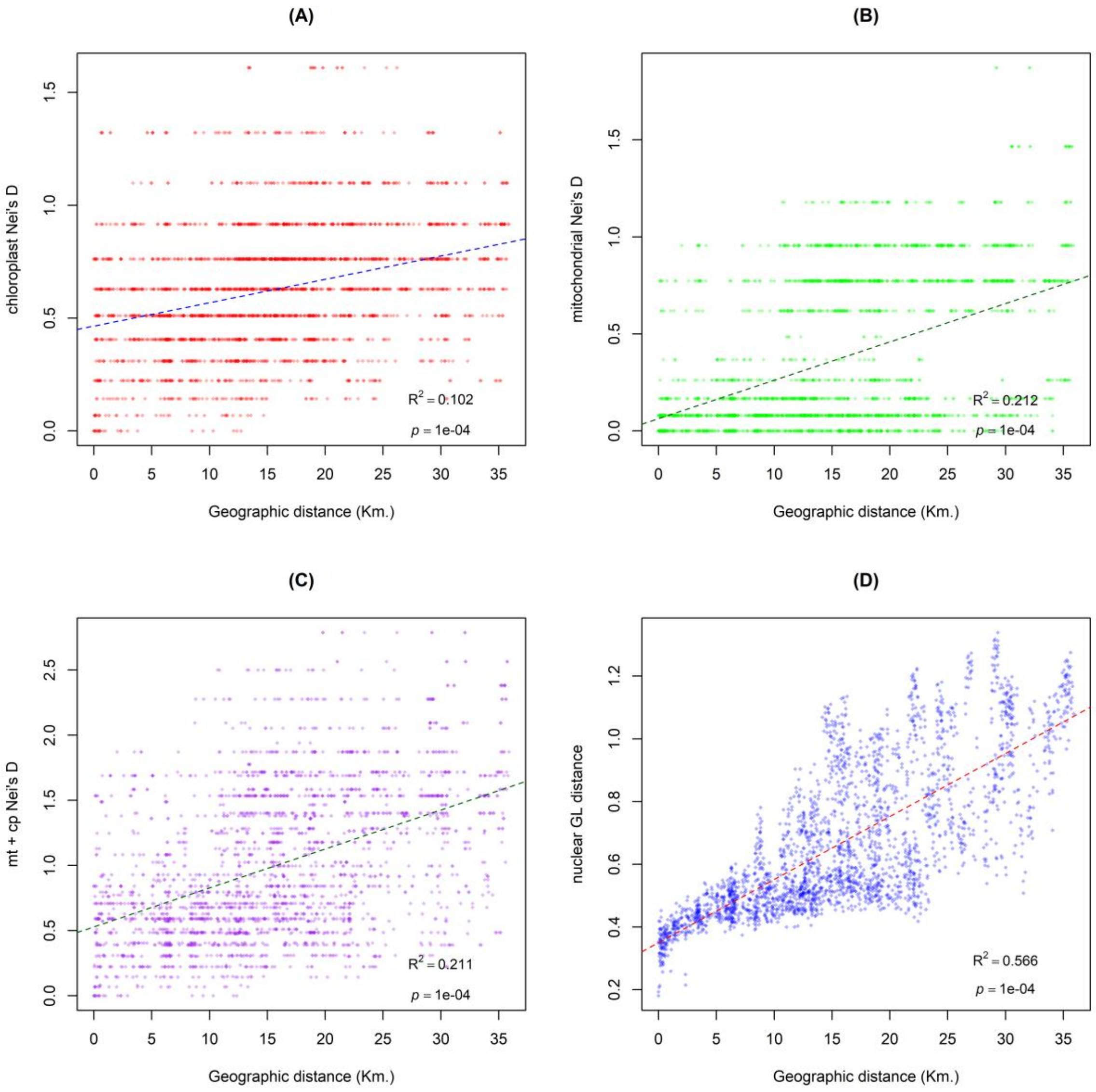


**Figure S15: Isolation by distance in *Noronhia spinifolia***

Graphic representation of the relationship between geographic and genetic distances (isolation by distance). The individual-based genetic distances estimated from chloroplast microsatellite data **(A),** mitochondrial data **(B)**, combined organellar DNA data **(C)**, and nuclear RAD data **(D)** and showing the highest relationship with geographic distance (Fig. S13) are represented against geographic distance. The linear model (dashed lines), the Mantel tests explained variance ($R^2$) and *p*-value (*p*) are shown to support the represented trends. The figure shows a positive relationship for both chloroplast, mitochondrial and nuclear data with Euclidean geographic distances suggesting that the gene flow may be impacted by geographic features. It further shows a stronger relationship for nuclear than for organellar DNA data.

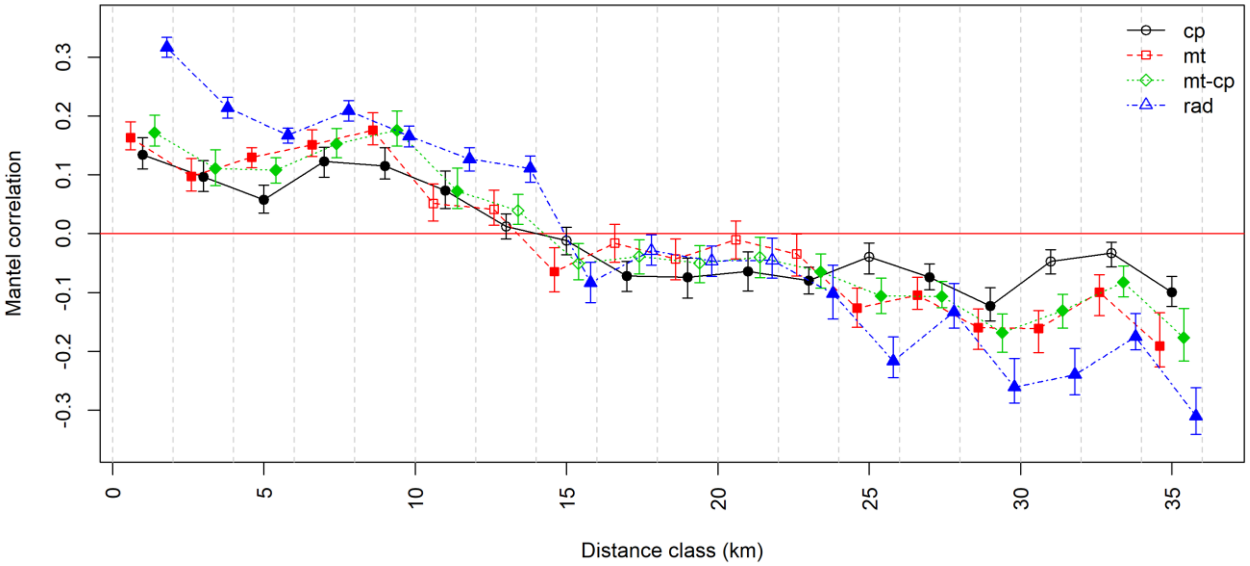


**Figure S16: Mantel correlogram of spatial correlation**

Mantel correlogram representation of the isolation by distance (IBD) relationship between individual-based genetic and geographic distances, estimated for 18 classes of 2 Km, with nuclear RAD (rad - PCA covariance distance), chloroplast (cp - Nei's *D*), mitochondrial (mt - Nei's *D*) and the combination of both mitochondrial and chloroplast distances (mt-cp - Nei's *D*). Significant values, over 1000 permutations, are represented by filled symbols.

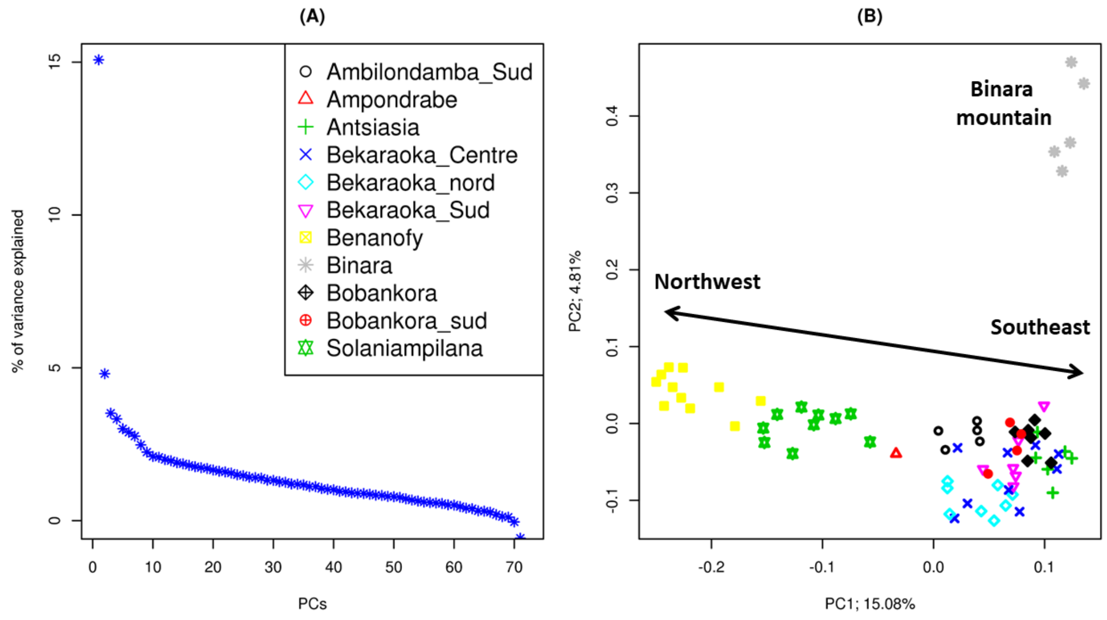


**Figure S17: Principal component analysis of nuclear genomic data of *Noronhia spinifolia***

Principal component analysis of *N. spinifolia* based on nuclear RAD-seq genotype likelihood, conducted in PCAngsd (Meisner & Albrechtsen, 2018). In **(A),** the proportion of variance explained by each of the principal components (PC) suggests that the two first components explain a notably large portion of the variance. In **(B)**, the representation of individual positions on these two components shows a strong and continuous cline from northwest to southeast along the first component axis that explains as much as ~15% of the variance. The second axis (explaining ~4.8% of the variance) clearly separates Binara from the other forests.

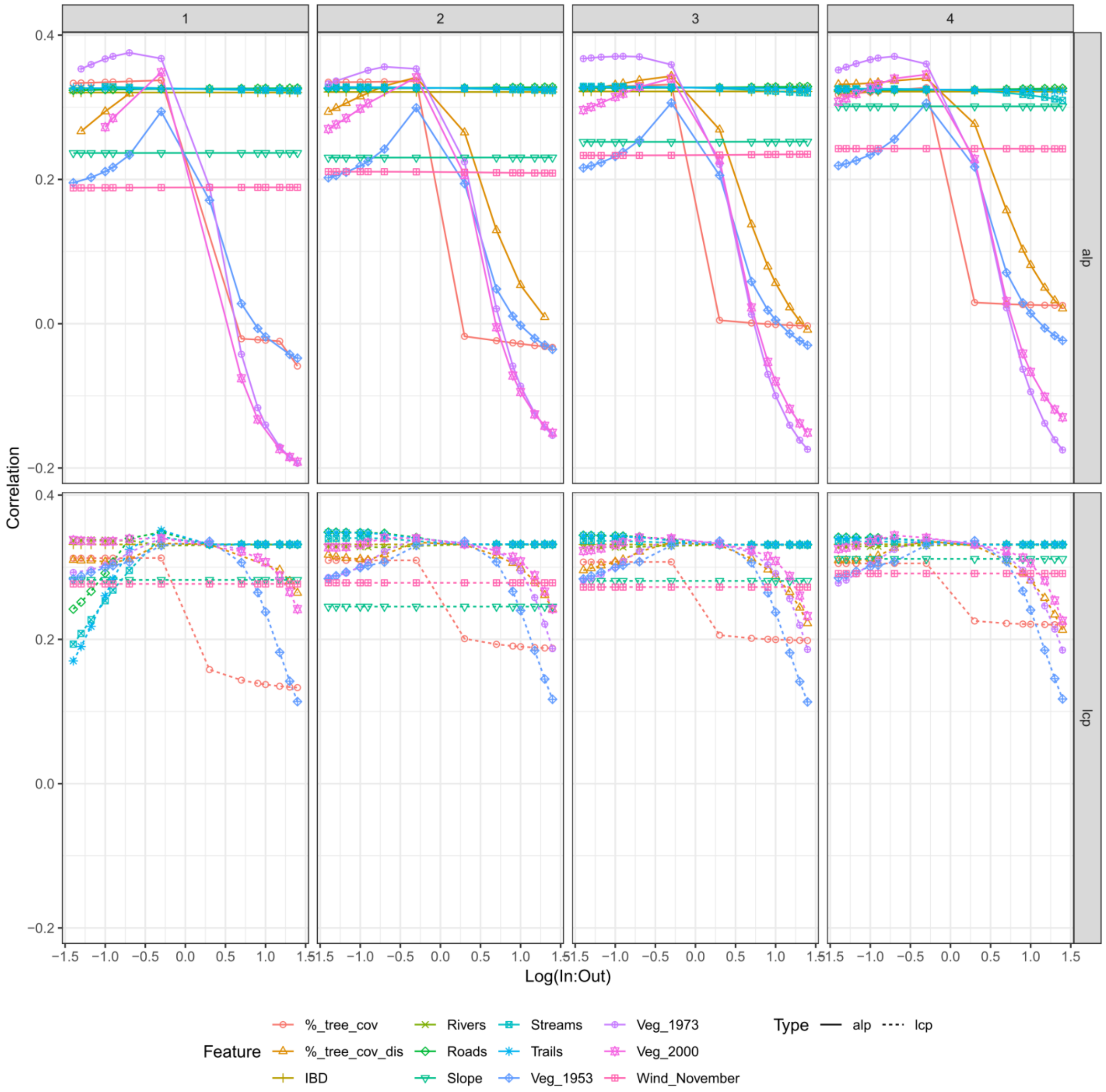


#### Figure S18: Univariate variable selection for chloroplast data

Representation of the landscape and genetic distances (isolation by resistance) univariate model fit. The explained variance (Correlation; *R*) of individual-based Mantel tests between landscape and genetic distances estimated from cpSSRs are represented as a function of the resistance values (x-axis: Log(in:out); the logarithm of the ratio between cost variable and cost non-variable; from high conductance on the left and to high resistance on the right of each panel; cf. Method S10), for varying granularity (pixel size 1-4, left to right panels), movement models (Type: alp = Circuit theory, upper panel; lcp = least cost path, lower panel), and landscape features (Feature). %_tree_cov and %_tree_cov_dis: Percent tree cover (Hansen et al., 2013) used as a continuous variable, or as a discrete variable with only values > 40%. IBD: Layer with the same cost in each cell therefore representing simple IBD and used as a null model. Veg_1953-1973-2000: forest covers (Vieilledent et al., 2018) from respective periods.

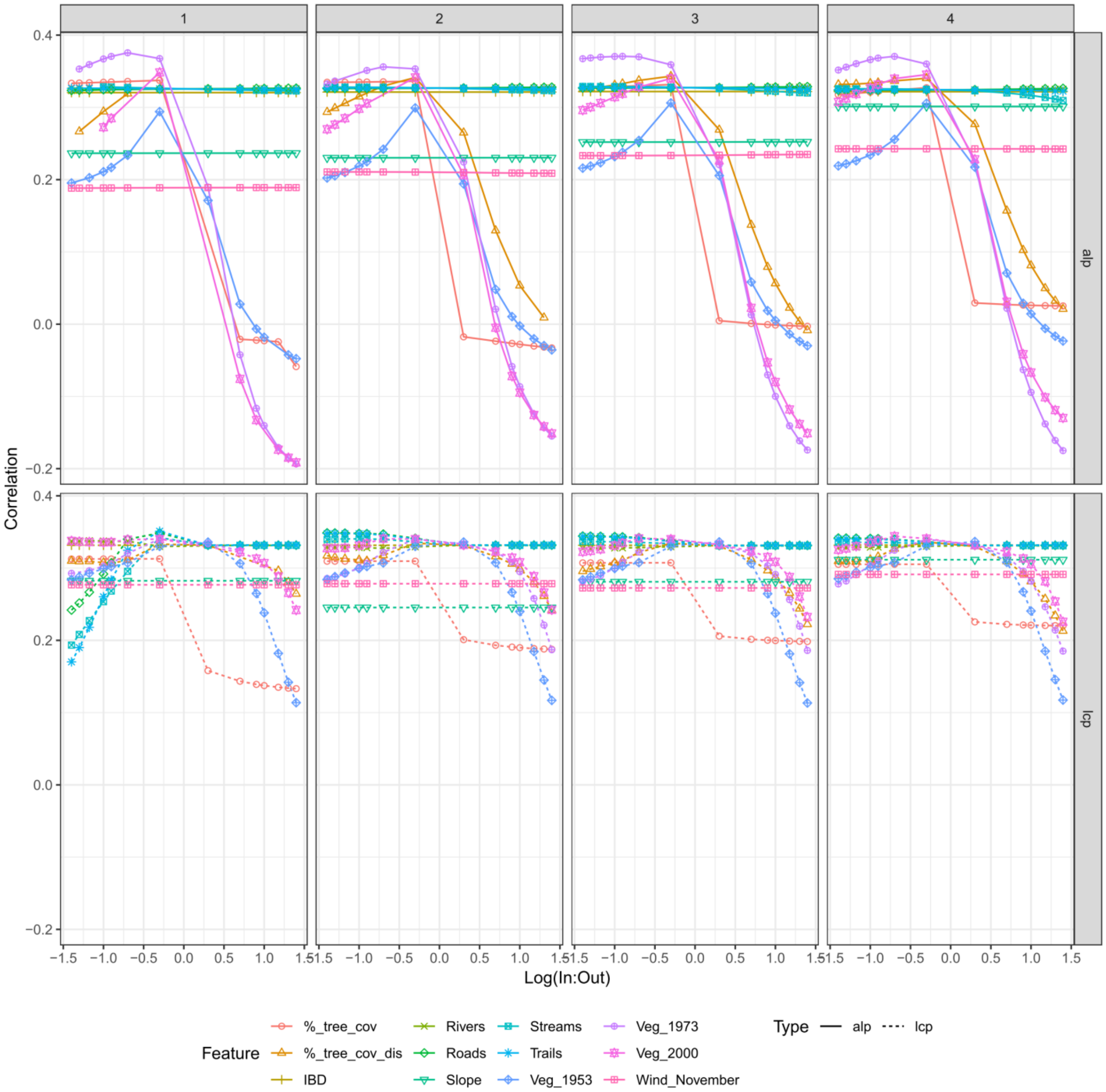


#### Figure S19: Univariate variable selection for mitochondrial data

Representation of the landscape and genetic distances (isolation by resistance) univariate model fit. The explained variance (Correlation; *R*) of individual-based Mantel tests between landscape and genetic distances estimated from mtRAD SNPs are represented as a function of the resistance values [x-axis: Log(in:out) = logarithm of the ratio between cost variable and cost non-variable on a log scale (from high conductance on the left to high resistance on the right of each panel); Method S10], for varying granularity (pixel size 1-4, left to right panels), movement models (Type: alp = Circuit theory, upper panel; lcp = least cost path, lower panel), and landscape features (Feature). %_tree_cov and %_tree_cov_dis: Percent tree cover (Hansen et al., 2013) used as a continuous variable, or as a discrete variable with only values > 40%. IBD: Layer with the same cost in each cell therefore representing simple IBD and used as a null model. Veg_1953-1973-2000: forest covers (Vieilledent et al., 2018) from respective periods.

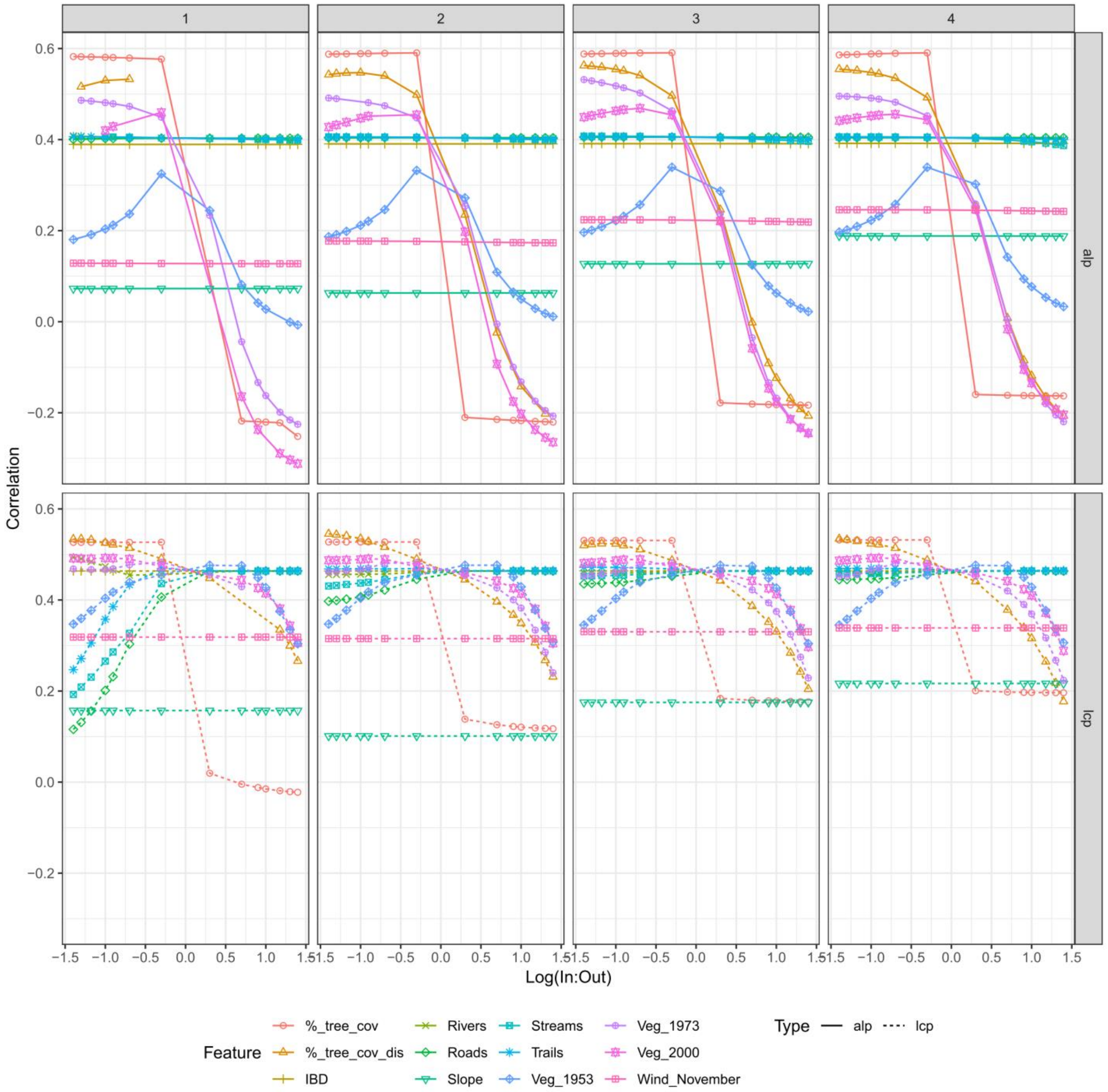


#### Figure S20: Univariate variable selection for organellar data

Representation of the landscape and genetic distances (isolation by resistance) univariate model fit. The explained variance (Correlation; *R*) of individual-based Mantel tests between landscape and genetic distances estimated from mtRAD SNPs and cpSSRs (Bruvo) are represented as a function of the resistance values [x-axis: Log(in:out) = logarithm of the ratio between cost variable and cost non-variable (from high conductance on the left to high resistance on the right of each panel); Method S10], for varying granularity (pixel size 1-4, left to right panels), movement models (Type: alp = Circuit theory, upper panel; lcp = least cost path, lower panel), and landscape features (Feature). %_tree_cov and %_tree_cov_dis: Percent tree cover (Hansen et al., 2013) used as a continuous variable, or as a discrete variable with only values > 40%. IBD: Layer with the same cost in each cell therefore representing simple IBD and used as a null model. Veg_1953-1973-2000: forest covers (Vieilledent et al., 2018) from respective periods.

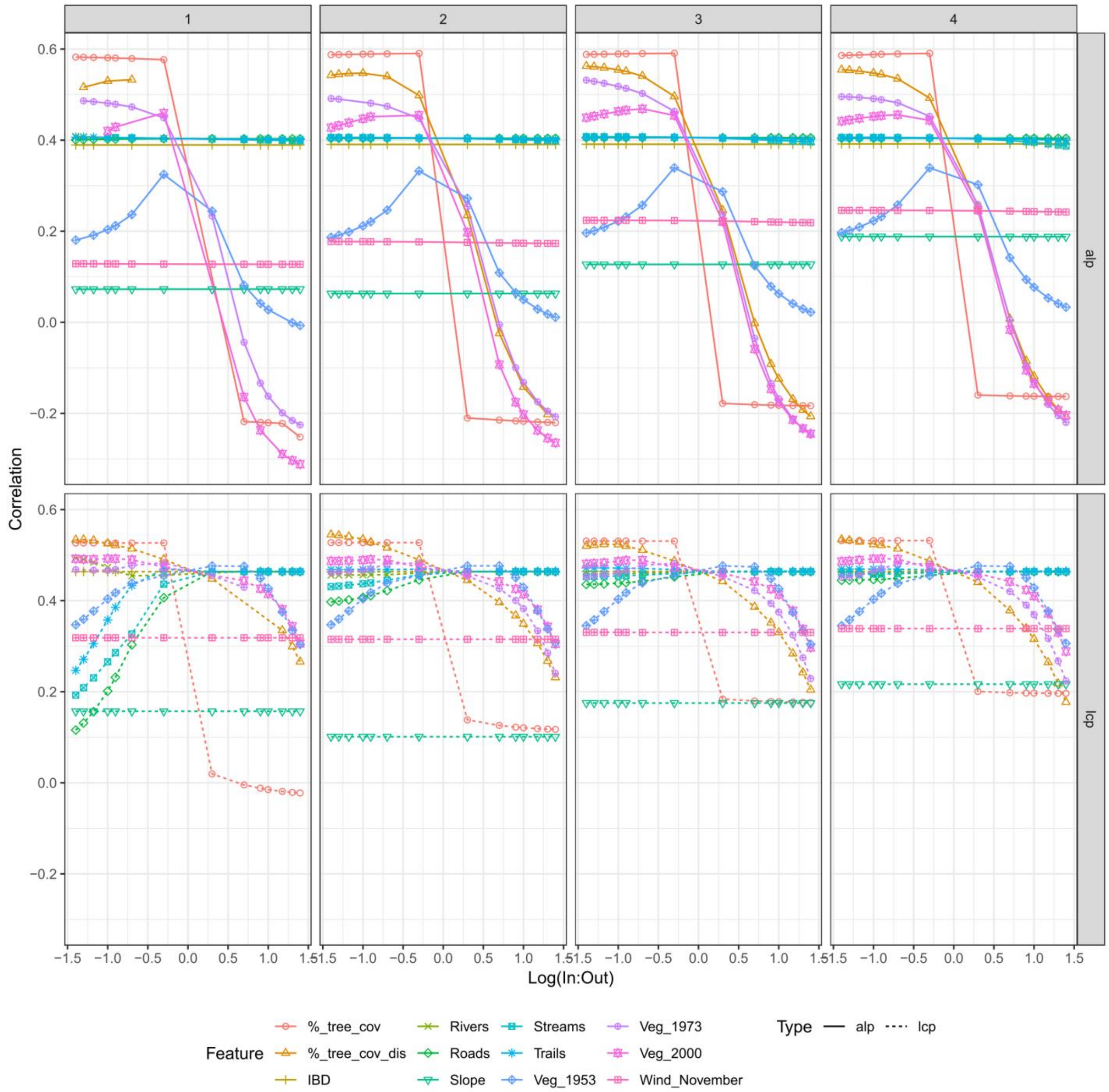


#### Figure S21: Univariate variable selection for nuclear data

Representation of the landscape and genetic distances (isolation by resistance) univariate model fit. The explained variance (Correlation; *R*) of individual-based Mantel tests between landscape and genetic distances estimated from nuclear data (GL.cov) are represented as a function of the resistance values [x-axis: Log(in:out) = logarithm of the ratio between cost variable and cost non-variable (from high conductance on the left to high resistance on the right of each panel); Method S10], for varying granularity (pixel size 1-4, left to right panels), movement models (Type: alp = Circuit theory, upper panel; lcp = least cost path, lower panel), and landscape features (Feature). %_tree_cov and %_tree_cov_dis: Percent tree cover (Hansen et al., 2013) used as a continuous variable, or as a discrete variable with only values > 40%. IBD: Layer with the same cost in each cell therefore representing simple IBD and used as a null model. Veg_1953-1973-2000: forest covers (Vieilledent et al., 2018) from respective periods.

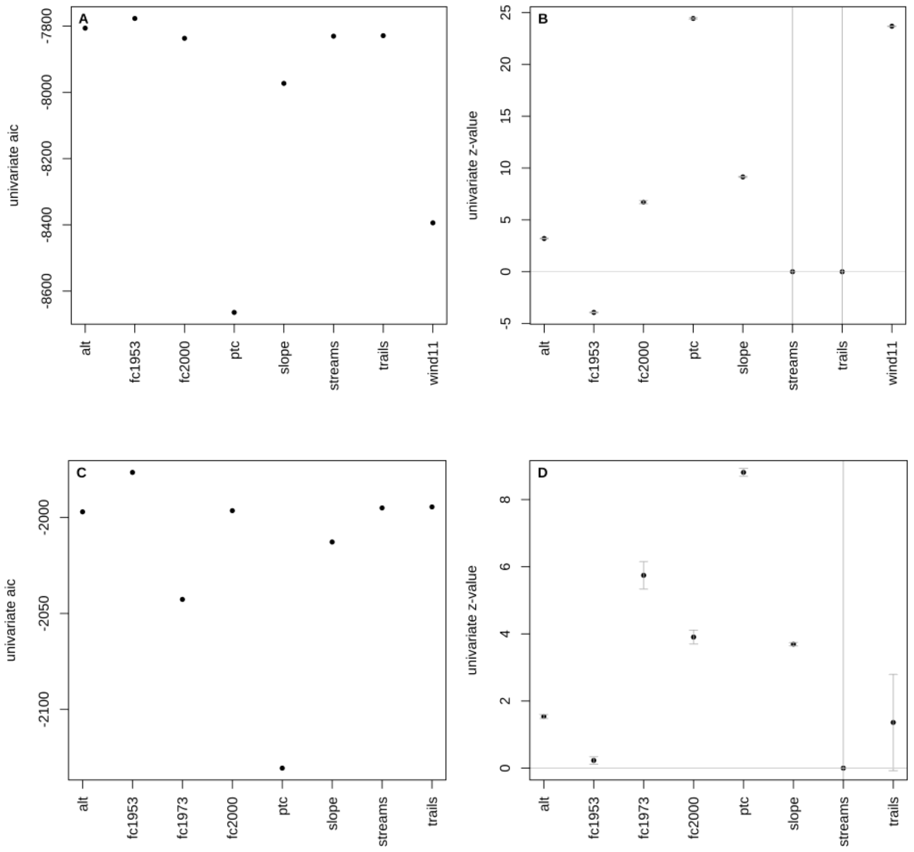


#### Figure S22: Univariate optimized association with genetic distances.

Representation of the univariate models AIC goodness of fit (**A, C**) and of the z-value (**B, D**, gray vertical bars: standard error) of the main retained landscape variable association with nuclear (**A-B**) and organellar (**C-D**) genetic distances, estimated in *radish* (Peterman & Pope, 2021) with a log-linear conductance and a MLPE measurement model. The percent tree cover (ptc), the slope (slope) and the wind speed in November (wind11) show the best fitting model (lowest AIC in **A**), and exhibit the strongest positive associations with nuclear genetic distances (z-value in **B**). In contrast, only the forest layers and the slope show good fitting model (lowest AIC in **C**), and exhibits the positive associations with organellar genetic distances (z-value in **D**). Streams and trails show large z-value standard error (gray vertical bars in C), not significantly different from zero (no association) and not always entirely represented within the area of the graphic. alt: altitude, fc1953: forest cover in 1953, fc1973: forest cover in 1973, fc2000: forest cover in 2000.

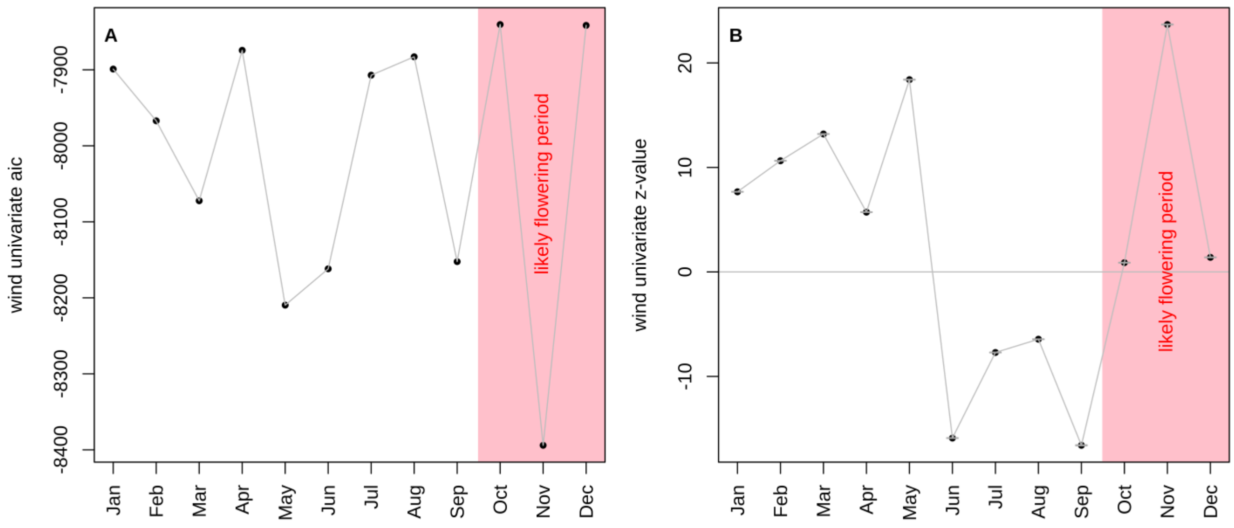


**Figure S23: Univariate monthly wind speed association with nuclear genetic distances.**

Representation of the AIC goodness of fit (**A**) and of the z-value (**B**) of monthly wind speed association with nuclear genetic distances (gray vertical bars: standard error), estimated in *radish* (Peterman & Pope, 2021) with a log-linear conductance and a MLPE measurement model. *Noronhia spinifolia* most likely flowering period identified from herbarium (Geneva and Paris) specimens' phenology is shaded in pink and includes the best fitting monthly wind speed model (November, lowest AIC in **A**), which exhibits the strongest positive association with nuclear genetic distances (z-value in **B**).

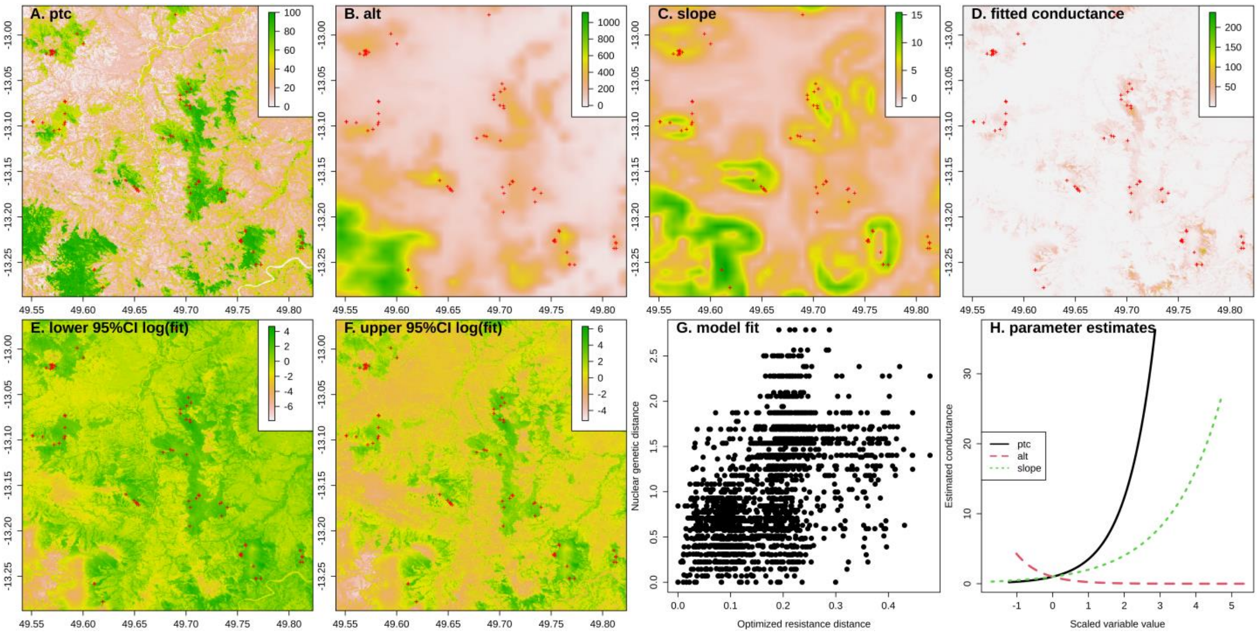


**Figure S24: Landscape contribution to organellar gene flow in *Noronhia spinifolia*.**

The figure shows the three landscape variables (**A-C**) likely contributing to *N. spinifolia* organellar gene flow (seeds dispersal), their fitted composite conductance (**D**) and associated confidence interval (**E-F**), the model fit relationship between the optimized resistance distance and the nuclear genetic (**G**), and the relative effects of the three retained variables (**H**). It illustrates a major conducting effect of the current forest cover (ptc: percent tree cover) on the connectivity of *N. spinifolia*, and it further shows a composite effect of the topography (positive effect of slope and negative effect of altitude [alt]).
